# DTX4 regulates neural progenitor transitions during cortical development

**DOI:** 10.1101/2025.04.10.648213

**Authors:** Elisa Pedersen, Sicheng Zhang, Xushuai Dong, Valeria Fernandez Vallone, Ekaterina Epifanova, Paul Moritz Willecke, Denis Lajkó, Theres Schaub, Mateusz Cyryl Ambrozkiewicz, Kathrin Textoris-Taube, Harald Stachelscheid, Marta Rosário

## Abstract

Neural progenitors drive cellular diversification in the neocortex. Prolongation of their proliferative capacity and increased diversity underpins the evolutionary expansion and morphological complexity of the human neocortex. Here, we investigate the mechanisms that regulate maintenance of the highly proliferative early neural progenitor subtypes and transition to subsequent progenitors of limited proliferative capacity during human and murine neocortical development. We identify DTX4, a Deltex family member, as an evolutionarily conserved molecular determinant of neural progenitor identity in the mammalian neocortex. DTX4 sustains the identity of radial glia, a highly proliferative progenitor subtype. Loss of DTX4, on the other hand, is a prerequisite for the generation of intermediate progenitors which possess low proliferative capacity. Perturbing DTX4 expression in human cerebral organoids and in the murine neocortex, alters progenitor composition, thereby inducing changes in neuronal diversity and cortical morphology. Mechanistically, we reveal that DTX4 controls progenitor identity by regulating the length of the cell cycle. Our findings underscore the critical role of cell cycle dynamics and of DTX4 in determining progenitor identity and thereby defining neuronal outcome during mammalian development.

**Highlights:**

● The Deltex family member, DTX4, is expressed in human and murine neuroepithelia and radial glial progenitors but is absent in intermediate progenitor cells.
● DTX4 maintains radial glia identity and prevents generation of intermediate progenitors during murine and human development.
● Disruption of DTX4 expression compromises neuronal composition and neocortical architecture
● DTX4 regulates progenitor identity by regulating cell cycle length

## Introduction

The neocortex owes its remarkable evolutionary expansion and complexity to the diversity and precise regulation of neural progenitor dynamics during development. Neocortical progenitors emerge in a temporally and spatially ordered sequence. Neuroepithelial cells (NECs), the earliest progenitors, initially undergo symmetric divisions to expand the progenitor pool before transitioning to asymmetric divisions, giving rise to radial glia cells (RGCs) around embryonic day (E) 10 in mice and gestational week (GW) 7 in humans (reviewed in ^1,2^). RGCs, the primary neural progenitors, are involved in neocortical expansion and neurogenesis but are also essential for radial migration and the laminar localization of neurons. During early development, RGCs proliferate symmetrically to further amplify the progenitor pool and later switch to asymmetric divisions to self-renew and generate either deep-layer (DL) excitatory neurons or intermediate progenitor cells (IPCs). IPCs are a division-limited progenitor population that in the murine neocortex contribute neurons to all cortical layers (reviewed in ^3^). Evolutionary expansion of the neocortex has been accompanied by increased progenitor diversity, including the emergence of outer RGCs (oRGCs), a progenitor subtype with high proliferative capacity which is scarce in lissencephalic animals^4^.

Regulation of progenitor cell cycle dynamics and mode of division is critical for determining the size, diversity, and organization of the neocortex. Disruptions in these processes can lead to severe neurodevelopmental disorders, such as microcephaly^5^. As corticogenesis proceeds, progenitor subtypes exhibit distinct cell cycle kinetics, with a progressive lengthening of the cell cycle correlating with the generation of more neuronal progeny^6–9^. However, the mechanisms underlying these temporal changes in cell cycle dynamics and their impact on progenitor maintenance and fate transitions remain poorly understood.

Here, we identify DTX4, a member of the Deltex family, as a key regulator of RGC maintenance and cell cycle progression during murine and human neocortical development. We demonstrate that DTX4 promotes the RGC progenitor state, and prevents transition to intermediate progenitors. DTX4 ensures rapid cell cycle progression and prolongs the RGC phase, thereby directing neural diversity and cortical morphology. Our findings uncover a previously unknown role for DTX4 in corticogenesis and provide new insights into the mechanisms linking cell cycle regulation to neural progenitor fate and cortical development.

## Results

### Early but not intermediate progenitor cells express DTX4

Neocortical size is determined by the extent of early neural progenitor proliferation. To investigate the mechanisms underlying neocortical expansion, we focused on early neural progenitors, whose proliferative capacity is a key determinant of neocortical size. We hypothesized that genes that are strongly expressed in neuroepithelial and radial glial (RGCs) progenitors but then downregulated upon transition to less proliferative intermediate progenitors (IPCs), may regulate this process. We focused on E3 ubiquitin ligases due to the strong association of neurodevelopmental disorders with disruption of these genes^10^. Analysis of differentially expressed E3 ubiquitin ligases using publicly available single cell RNA sequencing (scRNAseq) data from human cerebral organoids at D30, a timepoint where the first deep layer neurons are born^11^, led to the identification of DTX4, a member of the Deltex family of putative E3 ubiquitin ligases^12^. Deltex proteins have been implicated in the regulation of cell proliferation and differentiation in other tissues, but their function in the brain is unknown^12^. Of the Deltex members, *DTX1, DTX2, DTX3* and *DTX4* all show significant expression in human cerebral organoids at D30 (Fig, 1A, Supplementary Fig. 1). However, only *DTX4* is differentially expressed in RGCs (*SOX2^+^* population) compared to IPCs (*EOMES+* population) (Fig. 1A). Intriguingly, human *DTX4* shows a biphasic expression pattern, with high transcript levels in human RGCs, followed by sharp downregulation in IPCs, and subsequent re-expression in neurons (such as in *BCL11B^+^*deep layer) (Fig. 1A). DTX4 is strongly expressed in different RGC subtypes including oRGC (*HOPX^+^SOX2^+^*) and aRGC (*HOPX^-^SOX2^+^*) populations (Fig. 1A).

**Figure 1.**
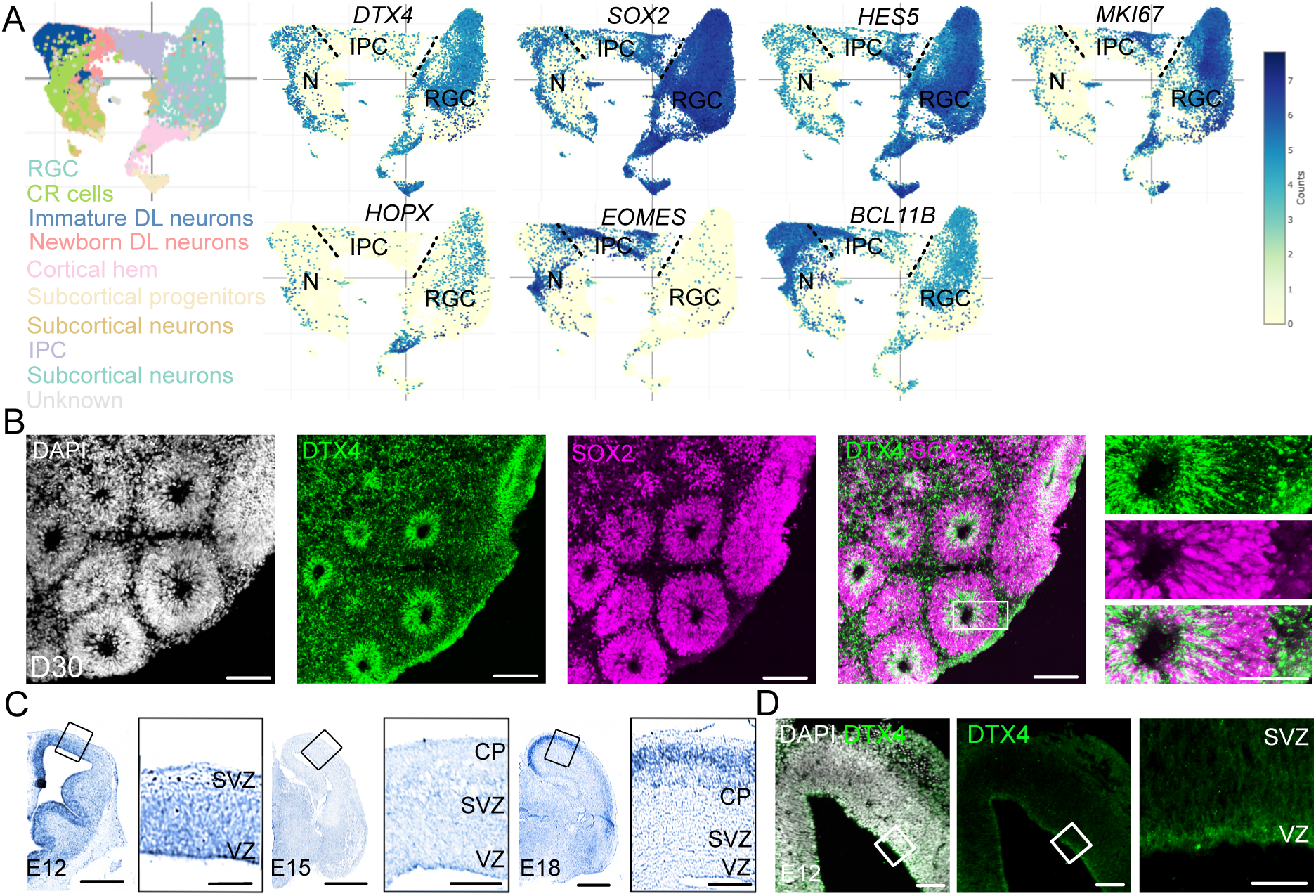
Early but not intermediate progenitor cells express DTX4 during human and murine neocortical development. **(A) DTX4 shows biphasic expression during human cortical development.** (A) scRNAseq data from D30 human cerebral organoids from Uzquiano *et al (*2022)^11^. UMAP representation of gene expression profiles for *DTX4* and representative marker genes for major cell types: RGC (*SOX2^+^*), NEC and RGCs (*HES5^+^*), proliferating cells (*MKI67^+^*) oRGCs (*HOPX^+^*), IPCs (*EOMES^+^*), immature and mature DL neurons (*BCL11B^+^*). RGC, radial glial cell; IPC, intermediate progenitors; N, immature and mature neurons. See also Supplementary Fig. S1. **(B) DTX4 is expressed in human RGCs during development.** DTX4 protein is expressed in human SOX2^+^ RGCs at D30. Immunofluorescent staining of D30 human cerebral organoids for DTX4 (green), SOX2 (magenta), and DAPI (grey). Scale bar = 200μm (overview), 50μm (magnified region). See also Supplementary Fig. S2. **(C) DTX4 is expressed at different stages of murine neocortical development.** ISH for DTX4 in mouse brains at E12, 15, and 18. Scale bar E12 = 400μm, magnified scale bar = 100μm. Scale bar E15 = 500μm, magnified scale bar = 150μm, Scale bar E18 = 800μm, magnified scale bar = 200μm. VZ, ventricular zone; SVZ, subventricular zone; CP, cortical plate. See also Supplementary Fig. S3-4 **(D) DTX4 protein is expressed in the VZ during early mouse cortical development**. E12 coronal mouse brain sections were stained for DTX4 (green) and DAPI (grey). Scale bar = 100μm (overview), 20μm (magnification).

To validate the expression pattern of DTX4 at the protein level, we performed immunofluorescence on human cerebral organoids using a specific DTX4 antibody (Fig. 1B; Supplementary Fig. S2A-B). At D30, we observed high levels of DTX4 protein in SOX2^+^ RGCs, particularly in RGCs lining the organoid lumen. As organoids matured, DTX4 expression in the ventricular zone (VZ) progressively decreased, consistent with the progressive decrease in RGCs numbers (Supplementary Fig. S2B). We also observed DTX4 expression in deep layer (DL, BCL-11B^+^) neurons in these older organoids (Supplementary Fig. S2B).

To further validate the biphasic expression of DTX4 we performed in situ hybridization (ISH) for DTX4 in the developing mouse brain (Fig. 1C). Consistent with our observations in human cerebral organoids, murine *DTX4* was highly expressed in the proliferative ventricular zone (VZ) at embryonic day 12 (E12), a region predominantly populated by RGCs at this stage. As development progressed, *DTX4* expression in the VZ and subventricular (SVZ) zone diminished, becoming nearly undetectable by E15. Strikingly, *DTX4* expression re-emerged during late gestation (E18), in the cortical plate (CP), coinciding with the accumulation of mature neurons in this layer at this stage (Fig. 1C). Examination of a publicly available scRNAseq dataset from the developing mouse neocortex^13^, verified high expression of *DTX4* in RGCs, downregulation in IPCs and subsequent re-expression in mature excitatory neurons (Supplementary Fig. S3). Immunostaining of E12 mouse brains further confirmed high expression of DTX4 protein in the VZ, indicating expression in RGCs (Fig. 1D).

Notably, DTX4 is the only Deltex family members to show biphasic expression during both murine and human cortical development (Supplementary Fig. S1, S4). DTX1 is primarily expressed in mature neurons, DTX2 and 3 are similarly expressed in all cell types and DTX3L is only weakly, if at all, expressed during cortical development (Supplementary Fig. S1, S4).

Together, our data indicate an evolutionary conserved, dynamic regulation of DTX4 expression during neocortical development highlighting its potential role in early cortical development.

### DTX4 maintains RGC progenitor identity and prevents IPC generation

Given that transition of RGC to IPC progenitors is accompanied by loss of DTX4 expression, we asked whether DTX4 regulates progenitor composition during corticogenesis. We downregulated DTX4 expression in RGCs by *in utero* electroporation (*IUE*) of a specific shRNA (shDTX4^mu^; Supplementary Fig. S5) into the mouse brain during early development (E12; Fig. 2A-D). Co-expression of GFP allowed the identification of electroporated cells. Two days later, brains were analysed for expression of Pax6, a marker for RGCs (Fig. 2A), and EOMES, a marker for IPCs (Fig. 2B). Downregulation of DTX4 (DTX4 KD) significantly reduced the number of Pax6^+^ RGCs from approximately 30% in the scrambled shRNA (shScr) control to around 10% in DTX4 KD (Fig. 2C). Loss of Pax6^+^ RGCs was accompanied by a parallel upregulation of the EOMES^+^ IPC progenitor population, indicating premature transition of RGCs to IPC generation upon DTX4 KD (Fig. 2D). Importantly, co-expression of the human DTX4 protein, insensitive to shDTX4^mu^ (Supplementary Fig. S5A), restored RGC:IPC balance, indicating both specificity and a cell-intrinsic role for DTX4 in maintaining RGC progenitor identity (Fig. 2A-D).

**Figure 2.**
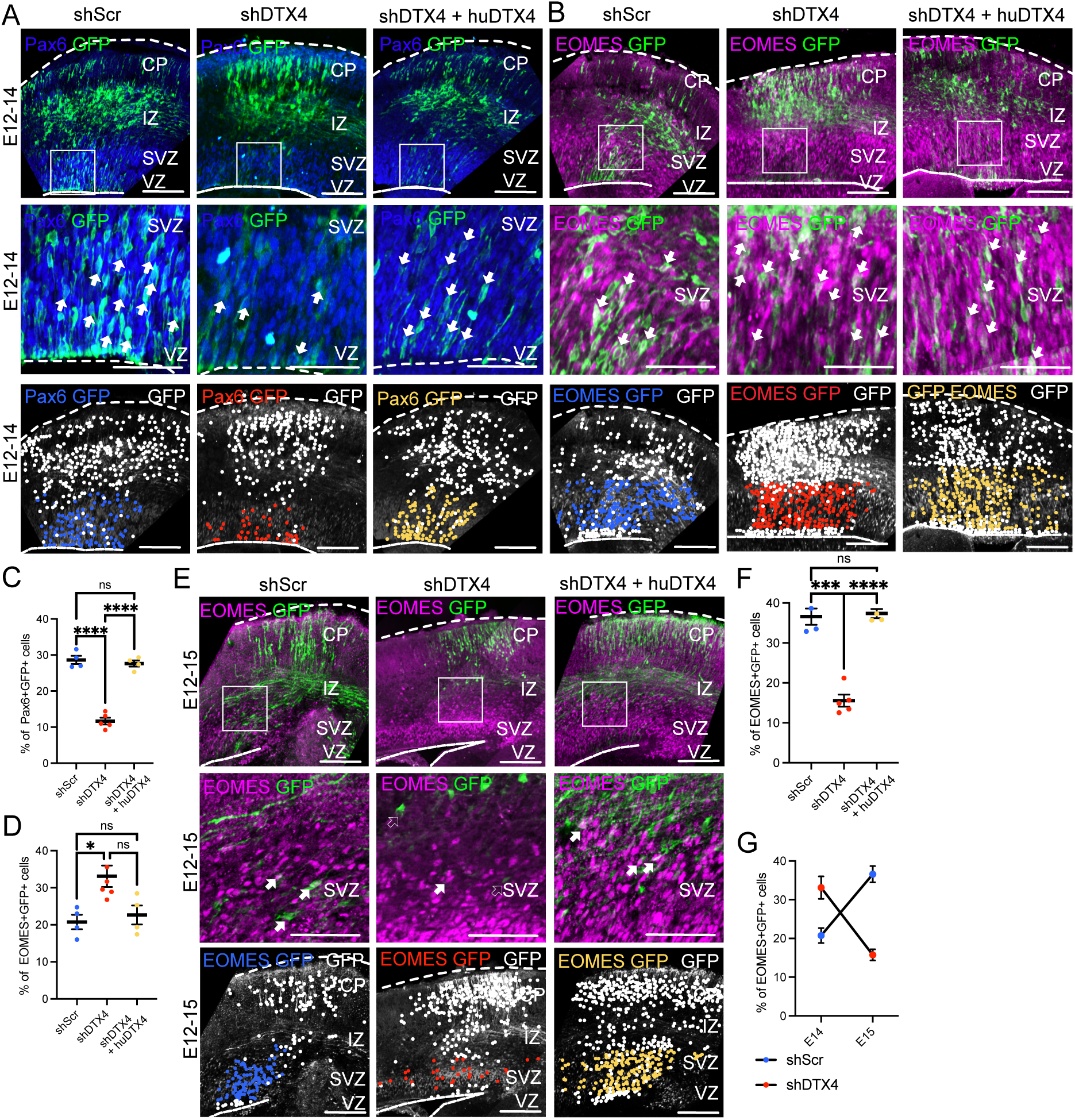
DTX4 maintains RGC progenitor identity and regulates progenitor transitions. **(A-D) DTX4 KD depletes RGCs and enhances generation of IPCs during murine corticogenesis.** Mice were *IUE* at E12 with a GFP expression construct and shScr, shDTX4 or shDTX4 + huDTX4 as indicated and stained at E14 for GFP (green) and either (A) Pax6 (blue) or (B) EOMES (magenta). Scale bar = 100μm. Pia and ventricular surfaces are marked with a dotted line. Magnified regions of the (A) VZ or (B) SVZ/IZ are shown below each representative image. Filled arrows indicate double positive cells. Scale bar = 50μm. The bottom row panels show the position of GFP^+^ soma (white dots), and double positive cells (coloured dots; Pax6^+^GFP^+^ or EOMES^+^GFP^+^) for each condition overlaying GFP staining (grey). Scale bar = 100μm. (C) Proportion of Pax6^+^ GFP^+^ and (D) GFP^+^ EOMES^+^ cells. N = 5 (shDTX4), 4 (shScr), 4 (shDTX4+huDTX4) animals. One-way Welch ANOVA with Dunnett’s correction. See also Supplementary Fig. S5 for validation of shRNAs used. **(E-G) DTX4 KD results in premature decrease in IPCs**. (E) Mice *IUE* at E12 with the indicated constructs were stained at E15 for GFP (green) and EOMES (magenta). Scale bar = 100μm. Magnified boxed regions are shown below. Filled arrows indicate double positive cells, empty arrows denote GFP^+^ EOMES^-^ cells. Scale bar = 50μm. (F) Proportion of GFP^+^ EOMES^+^ cells. N = 5 (shDTX4), 3 (shScr), 3 (shDTX4+huDTX4) animals. One-way Welch ANOVA with Dunnett’s correction. (G) Proportion of GFP^+^ EOMES^+^ cells at E14 and E15 N = 5 (shDTX4 at each timepoint), 3 (shScr at E15), 4 (shScr at E14). All data presented as individual points, mean and error bars with SEM.

To further trace the fate of DTX4 KD cells, we also analysed brains three days after *IUE* (Fig. 2E-F). At this later timepoint (E15), we observed a strong reduction in EOMES^+^ IPCs following DTX4 KD, consistent with the limited proliferative capacity of IPCs and their rapid subsequent differentiation into neurons (Fig. 2F). This delayed reduction in IPCs suggests that DTX4 KD effectively depletes the RGC progenitor pool, preventing further rounds of IPC generation (Fig. 2G). Together this data show that DTX4 is both necessary and sufficient to maintain RGC stemness. Furthermore, loss of DTX4 triggers premature transition to IPCs, ultimately disrupting the normal trajectory of corticogenesis.

### Initiation of neurogenesis and neuronal diversity are regulated by DTX4

Our findings indicate that loss of DTX4 causes premature generation of IPCs. To determine if this premature generation is associated with neurogenesis, we utilized a reporter mouse line for upper layer (UL) neurogenesis (Satb2^Cre/+^; ^14^). Primary cortical neurons from Satb2^Cre/+^ E12 embryonic brains were nucleofected with a dual-fluorescent Cre reporter construct (floxed mCherry-GFP^15^; Supplementary Fig. S6A). Induction of Satb2 expression, a marker and determinant for UL neurons^16^ results in Cre recombinase expression leading to excision of mCherry and expression of GFP. Flow cytometric analysis of GFP and mCherry expression 2 days after nucleofection, revealed increased production of Satb2^+^ (GFP^+^) neurons following DTX4 KD (Supplementary Fig. S6C-D), and decreased Satb2 neurogenesis following overexpression of DTX4 (DTX4 OE; Supplementary Fig. S6F-G). These results indicate that DTX4 regulates the initiation of UL neurogenesis.

Next, we manipulated DTX4 expression in RGCs by *IUE* of KD and OE constructs into wildtype mouse embryos at E12 and analysed the modified brains for neuronal progeny at E15 (Fig. 3A-E; Supplementary Fig. S7). Similar to our observations using the Satb2^Cre/+^ reporter line, DTX4 KD caused an enrichment of cells in the outer cortical layers, suggesting a critical role for DTX4 in determining the timing of neurogenesis (Fig. 3A-C). Furthermore, this shift in laminar position was restored upon expression of hDTX4 (Fig. 3B-C). To address whether the change in laminar position is associated with altered neurogenesis, we stained for both DL (BCL-11B^+^) and UL (Satb2^+^) neurons (Fig. 3A). DTX4 KD in RGCs resulted, three days later, in an increase in the proportion of both DL and UL neuronal subtypes (Fig. 3D-E), further indicating that loss of DTX4 drives progenitor transition and thereby premature neurogenesis. Expansion of Satb2^+^UL neurons was particularly striking (Fig. 3D). Satb2^+^ UL neurogenesis normally starts around E15^16^. In accordance at E15, less than 10% of control neurons expressed Satb2. In contrast, approximately 30% of DTX4 KD cells expressed Satb2, leading to distortion of the neuronal subtype composition towards UL cell fate (Satb2:BCL-11B ratio; Fig. 3F). Conversely, sustained expression of murine DTX4 had the opposite effect on neurogenesis (Supplementary Fig. S7A-F). At E15 DTX4-overexpressing cells occupied a much lower laminar position compared to control cells and both DL and UL neurogenesis were strongly suppressed, leading to a relative enrichment of DL neurons (Supplementary Fig. S7).

**Figure 3.**
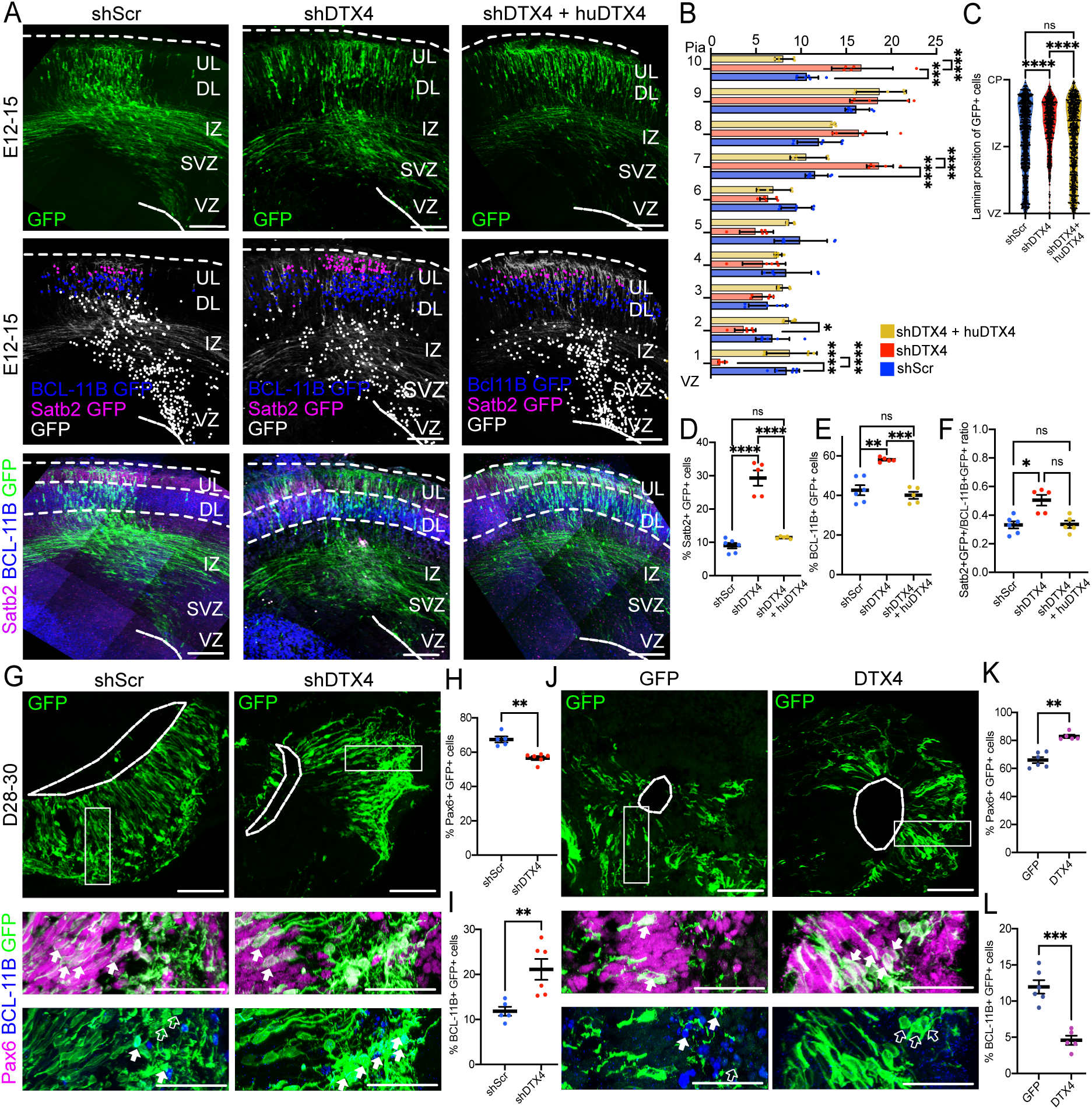
DTX4 regulates early neurogenesis. **(A-F) DTX4 KD accelerates neurogenesis in the murine cortex.** (A) E12 mice were *IUE* with a GFP expression construct and shScr, shDTX4 or shDTX4 + huDTX4 as indicated. Expression of GFP (green), BCL-11B (blue) and Satb2 (magenta) was analysed at E15. The position of GFP^+^ soma (white dots), and double positive cells (blue dots, BCL-11B^+^GFP^+^; magenta dots, Satb2^+^GFP^+^) for each representative image is shown below overlaying GFP staining (grey). Scale bar = 100μm. (B) Bin graph showing normalized laminar distribution of GFP^+^ across the cortex. N = 5 (shDTX4), 6 (shScr), 5 (shDTX4+huDTX4) animals. One-way ANOVA Tukey Multiple Comparison Test. (C) Laminar position of GFP^+^ cells. N = 1288 (shDTX4), 1091 (shScr), 1456 (shDTX4+huDTX4) cells from respectively 5, 6, 5 animals. Non-parametric Kruskal-Wallis test. See also Supplementary Fig. S6-7. (D) Proportion of Satb2^+^GFP^+^ cells. One-way ANOVA Tukey Multiple Comparison Test. (E) Proportion of BCL-11B^+^GFP^+^ cells. One-way Brown-Forsythe ANOVA with Dunnett’s Multiple Comparison. (F) Ratio of UL (Satb2^+^ GFP^+)^ to DL (BCL-11B^+^GFP^+^) neurons. Non-parametric Kruskal-Wallis test with Dunńs Multiple Comparison Test. N = 5 (shDTX4), 6 (shScr), 5 (shDTX4+huDTX4) animals in all cases. **(G-K) DTX4 controls RGCs and onset of neurogenesis in human cerebral organoids.** Human cerebral organoids were modified by *IOE* of (G-I) shScr or shDTX4^hu^ or (J-L) GFP or DTX4^hu^ as indicated and analysed at D30 for expression of Pax6 (magenta), BCL-11B (blue), and GFP (green) expression. Magnified regions are indicated by a box. Filled arrows indicate double positive cells, empty arrows indicate single positive GFP^+^ cells. Scale bar = 100μm (overview), 50μm (magnified region). Proportion of Pax6^+^GFP^+^ for (H) shScr and shDTX4^hu^ and (I) GFP and DTX4^hu^ conditions. Proportion of BCL-11B^+^ GFP^+^ cells for (J) shScr and shDTX4^hu^ and (K) GFP and DTX4^hu^ conditions. N = 6 organoids per condition. Kolmogorov-Smirnov test. All data presented as individual points, mean and error bars with SEM.

Together this data show that DTX4 prolongs RGC identity, preventing transition to IPC generation and subsequent production of their predominantly UL neuronal progeny. This is consistent with a potential contribution of DTX4 in the prolongation of early progenitor states associated with the evolutionary expansion of the neocortex. Given the differences in neural progenitor subtypes between murine and human corticogenesis, we investigated whether the role of DTX4 in sustaining RGC identity is evolutionarily conserved. To assess the functional conservation of DTX4, we downregulated DTX4 expression in human cerebral organoids via *in organoid* electroporation (*IOE*) of a shRNA targeting human DTX4 (shDTX4^hu^; Supplementary Fig. S5B). *IOE* was performed at D28, and human cerebral organoids were analysed two days later (D30) coinciding with the birth of the first DL neurons (Fig. 3G-I). Indeed, in control *IOE*, approximately 70% of GFP^+^ cells still expressed the RGC marker, Pax6, and only around 10% had started expressing the DL marker, BCL-11B by D30. Similar to what we observed in the mouse brain, hDTX4 KD in organoids caused depletion of RGCs and premature neurogenesis, as seen by the decrease in Pax6^+^ cells (Fig. 3H) and accompanying increase in BCL-11B^+^ DL neurons (Fig. 3I). Sustained expression of human DTX4 in human cerebral organoids, on the other hand, increased the number of RGCs and prevented neurogenesis, demonstrating that DTX4 is sufficient to prolong self-renewal of RGCs at the cost of neurogenic divisions (Fig. 3J-L). These findings underscore the evolutionary conservation of DTX4’s role in prolonging RGC identity and stemness during neocortical development.

### Downregulation of DTX4 in RGCs has enduring consequences for neural diversity and cortical organisation

To determine whether the neurogenic alterations induced by premature DTX4 downregulation have lasting consequences for neocortical development, we downregulated DTX4 expression in the mouse brain at E12 and analysed the outcomes at postnatal day 21 (P21) (Fig. 4). Control experiments with shScr and a GFP expressing construct produced no observable alterations in cortical structure or neuronal diversity. In stark contrast, DTX4 KD induced profound cortical malformations in all animals analysed (Fig. 4A-B). All DTX4 KD animals showed deep invaginations of the neocortex involving inwards folding of the cortical plate. These invaginations were consistently localized immediately medial to the *IUE*-targeted region. Importantly, the pia and the underlying white matter were not altered, indicating that invaginations did not result from *IUE*-induced cortical damage (Fig. 4A). In addition, 70% of animals showed heterotopias of variable size, that contain few GFP^+^ cells and were mostly composed of Cux2^+^ UL neurons, in the white matter of the electroporated hemisphere (Fig. 4C). *IUE* of a different, less efficient, shRNA against DTX4 had the same phenotypic outcome (shDTX4^mu2^; Supplementary Fig. S8A-B). Although we cannot definitively conclude whether total neuronal output is altered by DTX4 KD, distortion of the neocortical structure and the appearance of heterotopias are consistent with premature depletion of RGCs and associated disruption in the radial migration of later-born UL neurons. Electroporation of shDTX4^mu^ at E14, a timepoint where RGCs and thus also DTX4 expression is much reduced, had no effect on postnatal cortical morphology (Supplementary Fig. S9), underscoring the specificity of DTX4’s role in early progenitor function.

**Figure 4.**
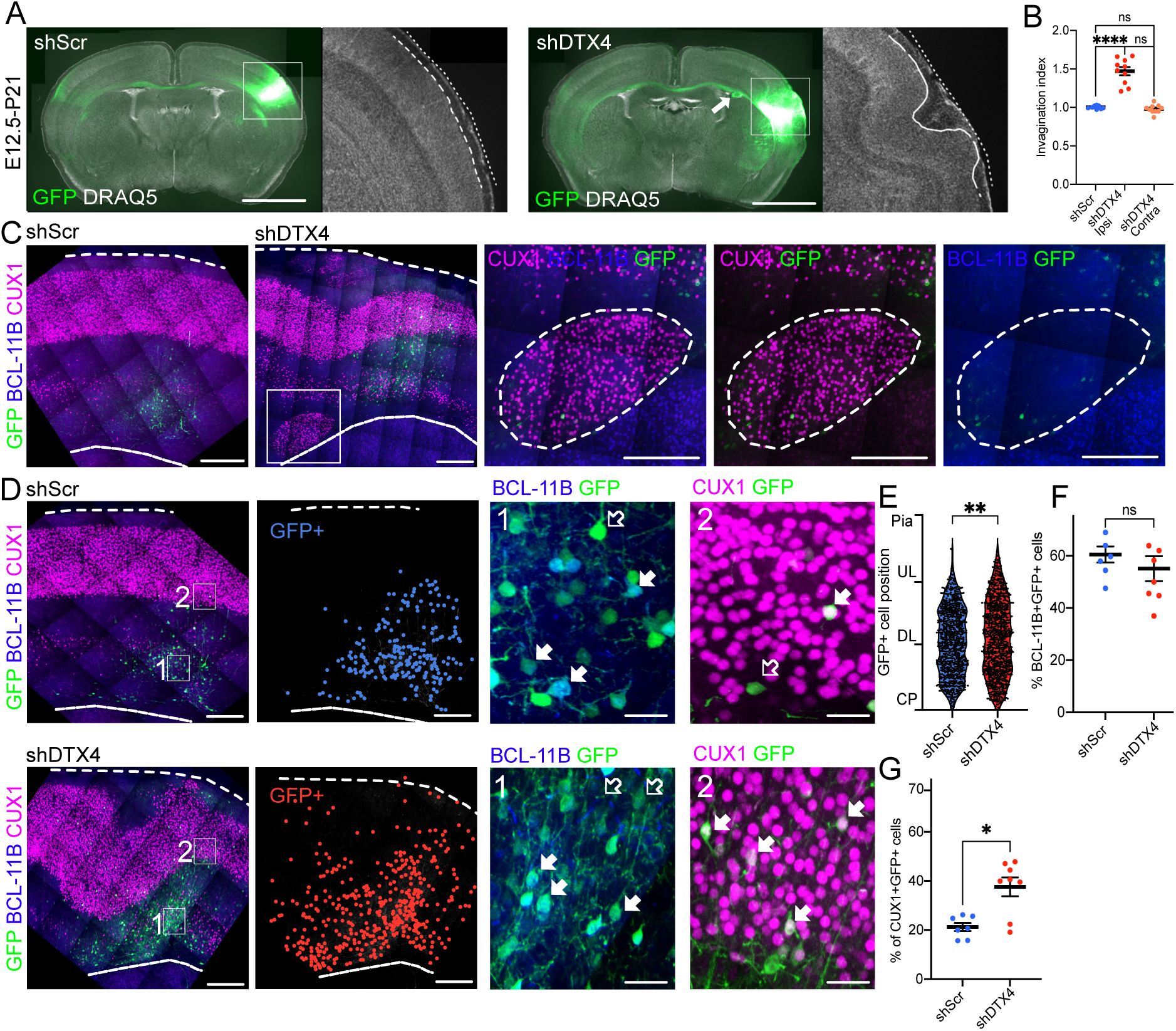
DTX4 function in RGCs has enduring consequences for neural diversity and cortical organisation. **(A-G) *IUE* of shScr or shDTX4 was carried out at E12 and animals were analysed at P21. (A-C) DTX4 regulates cortical morphology during early neurogenesis.** (A) Coronal sections of representative *IUE* brains stained for GFP (green) and DRAQ5 (grey). Arrow marks heterotopia as observed in 7/10 shDTX4 animals. Scale bar = 500μm. (B) Invagination index for *IUE* ispsilateral (ipsi) and contralateral (contra) hemispheres. N= 8 (shScr), 10 (shDTX4) animals. Normality evaluated using Shapiro-Wilk followed by one-way ANOVA with Tukey’s multiple comparisons test. N= 8 shScr, 10 shDTX4. (C) Staining for the UL marker, Cux1 (magenta), DL marker, BCL-11B, (blue), GFP (green) and DAPI (white) Dotted lines mark pia and ventricular surfaces. Scale bar = 200μm (overview) Magnification of boxed heterotopia is shown on the right. Scale bar = 50μm. **(D-G) Postnatal neuronal diversity is controlled by DTX4.** (D) Neuronal fate and laminar distribution were analysed at P21 by staining for BCL-11B (blue), Cux1 (magenta), GFP (green) and DAPI (grey). The position of GFP^+^ soma is marked by blue/red dots to the right of the merged image. Dotted lines mark pia and ventricular surfaces. Scale bar = 200μm. Magnified regions are boxed and numbered. Filled arrows indicate selected double positive cells and empty arrows mark single positive GFP^+^ cells. Scale bar = 25μm. (E) Laminar distribution of GFP^+^ cells with respect to ventricular surface. N = 825 shScr, 937 shDTX4 cells from 7 shScr, 8 shDTX4 animals. Kolmogorov-Smirnov test. (F) Proportion of BCL-11B^+^GFP^+^ cells. N = 7 (shScr), 8 (shDTX4) animals. Two tailed Welch’s t-test. (G) Proportion of Cux1^+^ GFP^+^ cells. N = 7 (shScr, 8 shDTX4 animals. Kolmogorov-Smirnov test. See also Supplementary Fig. S8-10. All data presented as individual points, mean and error bars with SEM.

Consistent with our findings during embryonic stages, analysis of the postnatal neuronal fate of cells where we downregulated DTX4 expression at E12, revealed a higher laminar position, increased proportion of Cux1^+^ UL neurons and a trend towards a decrease in the proportion of BCL-11B^+^ DL neurons (Fig. 4D-G). This was accompanied by a relative increase in UL neuron-derived midline-crossing GFP^+^ callosal axons suggesting normal neuronal differentiation of DTX4 KD neurons (Supplementary Fig. S10).

Collectively, these findings support a model in which DTX4 plays a critical role in maintaining the RGC identity and preventing transition to IPCs and neurogenesis. Disruption of this function leads to profound and lasting defects in neural composition and cortical structure, highlighting the importance of DTX4 during early corticogenesis.

### DTX4 is a key regulator of cell cycle dynamics

Cortical malformations, including abnormalities in gyrification, heterotopias, and microcephaly, are often linked to dysfunction or loss of RGCs. These disruptions lead to alterations in neuronal number, diversity, and laminar positioning^17^ similar to those observed upon downregulation of DTX4.

To assess if DTX4 influences the proliferative capacity of progenitors, we downregulated DTX4 in the mouse brain by *IUE* of shDTX4^mu^ at E12. 36h after *IUE* we exposed the embryos to a 2h pulse of the thymidine analogue 5-ethynyl-2’-deoxyuridine (EdU) to label cells in S phase (Fig. 5A-C). Remarkably, just 38h following *IUE*, DTX4-downregulated cells already occupy a higher laminar position (Fig. 5A-B), indicating very rapid consequences to changes in DTX4 expression for neurogenesis. DTX4 KD was also caused a reduction in EdU^+^ cells, indicating that DTX4 promotes RGC proliferation (Fig. 5A-C). Both laminar position and EdU incorporation were restored by expression of hDTX4 (Fig. 5A-C). Surprisingly, the proportion of Ki67^+^ cells, denoting the total proliferative population^18^, was not significantly altered by DTX4 KD during this short period (Supplementary Fig. S11). Sustained expression of DTX4 at E12 on the other hand, promoted EdU incorporation, also without in this short period altering the number of total dividing (Ki67^+^) cells (Supplementary Fig. S12). These data indicate that DTX4 is both necessary and sufficient to regulate RGC proliferation and suggests that this likely involves a change in cell cycle dynamics.

**Figure 5.**
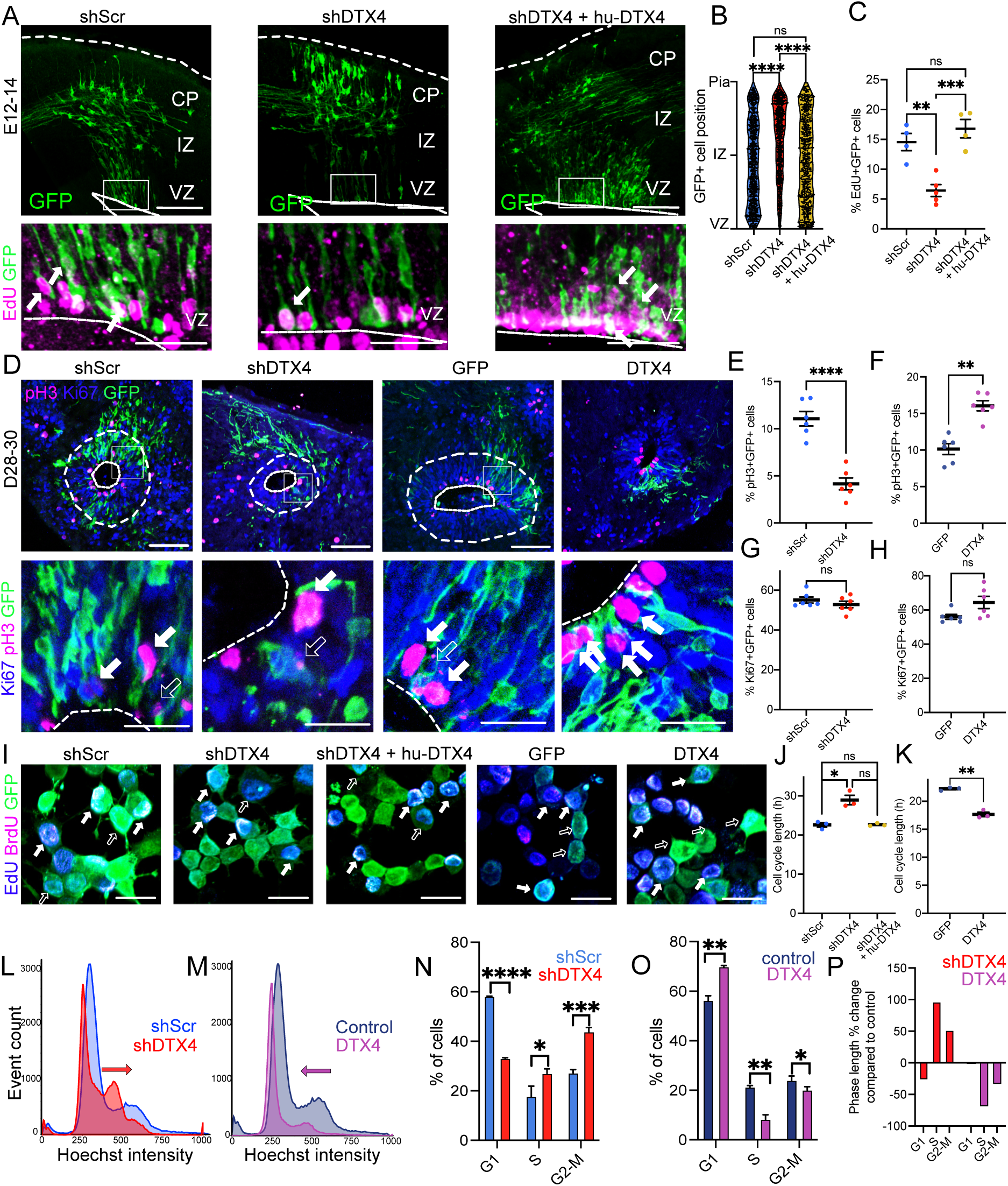
DTX4 is a key regulator of cell cycle dynamics in the neocortex. **(A-C) DTX4 regulates proliferation during murine development.** E12 mice were *IUE* with a GFP expression construct and shScr, shDTX4 or shDTX4+huDTX4 as indicated. Embryos were exposed to a 2h EdU pulse 36h after *IUE* and analysed at E14 for expression of GFP (green) and incorporation of EdU (magenta). (A) Representative images of *IUE* animals. Scale bar = 100μm. Magnified boxed areas are shown below. Arrows indicate double positive cells. Scale bar = 50μm. N = 5 (shScr), 4 (shDTX4), 4 (shDTX4+huDTX4) embryos. See also Supplementary Fig. S11-12. (B) Laminar distribution of GFP^+^ cells with respect to the ventricular surface. N= 823 (shScr), 892 (shDTX4), 715 (shDTX4+huDTX4) cells, respectively from 5, 4, 4 embryos. Non-parametric Kruskal-Wallis test. (C) Proportion of EdU^+^ GFP^+^ (S-phase) cells. N = 5 (shScr), 4 (shDTX4), 4 (shDTX4+huDTX4) embryos. One-way ANOVA with Dunn’s correction. **(D-H) DTX4 controls cell cycle dynamics in human cerebral organoids.** (D) Human cerebral organoids were *IOE* at D28 with shScr, shDTX4, GFP, or DTX4 as indicated and analysed at D30 for expression of GFP (green), DAPI (grey), pH3 (magenta, M-phase) and Ki67 (blue, proliferating cells). Scale bar = 100μm. Magnified areas are boxed. Scale bar = 25μm. Filled arrows indicate triple positive cells and empty arrows GFP^+^ cells. (E) Proportion of M phase (pH3^+^GFP^+^) cells in shScr and shDTX4 conditions. Two-tailed Welch’s T-Test. (F) Proportion of M phase (pH3^+^GFP^+^) cells in GFP and DTX4 conditions. Non-parametric Kolmogorov-Smirnov test. (G) Proportion of proliferating (Ki67^+^GFP^+^) cells in shScr and shDTX4. Non-parametric Kolmogorov-Smirnov test. (H) Proportion of proliferating (Ki67^+^GFP^+^) cells in GFP and DTX4. Two-tailed Welch’s T-Test. N = 6 organoids per condition in all cases. **(I-K) DTX4 regulates cell cycle duration.** (I) Neuro-2a cells were transfected with a GFP expression construct and either shScr, shDTX4, shDTX4 + huDTX4, GFP alone, or DTX4 as indicated. Cells were exposed to a 1.5 h spaced EdU-BrdU pulse 24h after transfection and analysed for expression of GFP (green), incorporation of EdU (blue) and BrdU (magenta) and expression of Ki67 (Supplementary Fig. S13) to determine the total proliferating population. Scale bar = 20μm. (J-K) Cell cycle length (Tc) for (J) DTX4-KD and (K) DTX4-OE conditions was calculated using Tc (h) = P × Ti /L where Ti = time between injections, P = total number of proliferating cells, L (leaving fraction) = number of BrdU^-^ EdU^+^ cells as previously described ^20^. N = 3 independent transfections per condition (shScr N1= 442, N2= 429, N3= 303, shDTX4 N1= 404, N2= 555, N3= 483, shDTX4+hu-DTX4 N1= 405, N2= 557, N3= 280, GFP N1= 414, N2= 474, N3= 390, DTX4 N1= 523, N2= 290, N3= 411). (J) Brown-Forsythe ANOVA test. (K) Two-tailed Welch’s T-Test. **(L-P) DTX4 regulates cell cycle dynamics**. (L-M) Representative cell cycle profiles for Hoechst 33342-stained Neuro-2a cells transfected 25h previously with a GFP expression construct and either (L) shScr (blue), shDTX4 (red) or (M) GFP (grey), DTX4 (magenta). (N-O) Proportion of cells in G1, S and G2/M phase following (N) DTX4 KD or (O) DTX4 OE. N = 3 (500 000 cells each). Two-tailed Welch’s T-Test. (P) Fold change in the duration of the different cell cycle phases upon DTX4 KD or OE. See also Supplementary Fig. S14. All data are represented as individual points, mean and error bars with SEM.

We extended our findings to human cerebral organoids, and either downregulated or overexpressed DTX4 at D28 and analysed the modified organoids 2 days later for the proportion of total proliferating cells (Ki67^+^) and also the proportion of cells found in M-phase by staining for the M-phase marker, phospho-Ser10 histone H3 (pH3; Fig. 5D-H). Similar to our observations in the murine neocortex, DTX4 KD and DTX4-OE altered cell cycle dynamics without affecting overall proliferation (Fig. 5D-H). Specifically, DTX4 KD strongly reduced the proportion of cells in M-phase (pH3^+^ cells; Fig. 5E) without affecting the overall proliferation in this short time period (Ki67^+^ cells, Fig. 5G). Inversely, overexpression of human DTX4 in organoids led to an increase in the proportion of M-phase cells (Fig. 5F) without altering overall proliferation (Fig. 5H). These results emphasize a conserved role for DTX4 in regulating the dynamics of neural progenitor proliferation, rather than cell cycle exit. The data is consistent with the impact of DTX4 on the composition of progenitor populations and neurogenic outcomes in both murine and human systems.

To further dissect the role of DTX4 in cell cycle dynamics, we measured the effect of DTX4 on the length of the cell cycle in asynchronous double pulse-chase experiments using thymidine analogues EdU and 5-bromo-2’-deoxyuridine (BrdU) in Neuro-2A cells, a highly proliferative murine cell line that endogenously expresses DTX4^19^. Staining for GFP, EdU and BrdU were used to determine the proportion of cells in S-phase and those that left S-phase in the 1.5h period between EdU and BrdU pulses, while staining for Ki67 was used to determine the total proliferative population (Supplementary Fig. S13). Cell cycle length was calculated from these values as previously described^20^. Control transfected cells showed a cell cycle length of approximately 22.5 hrs (Fig. 5I-K, Supplementary Fig. S13). Remarkably, DTX4 KD caused a substantial lengthening of the cell cycle (to 28.96 hrs), while DTX4 OE induced a marked shortening of the cell cycle (to 17.65 hrs; Fig. 5I, J) demonstrating that DTX4 has a direct and powerful impact on the duration of the cell cycle.

To further investigate the role of DTX4 in controlling cell cycle dynamics, we analysed the cell cycle profiles of Neuro-2A cells, 25 hrs (roughly one cell cycle length) after DTX4 KD or OE by FACS sorting of Hoechst 33342-stained cells (Fig. 5L-P). DTX4 KD induced a shift in the cell cycle profile characterised by accumulation of cells in S and G2/M, while DTX4 OE skewed the cell cycle towards G1 phase (Fig. 5L-P; Supplementary Fig. S14). These shifts show that DTX4 does not uniformly alter cell cycle length. Calculations based on cell cycle length (Fig. 5I-K) and cell cycle phase distributions (Fig. 5N-O) indicate that DTX4 has the greatest impact on S phase (Fig. 5P). Taken together these findings indicate that DTX4 maintains RGC stemness through regulation of cell cycle dynamics.

## Discussion

The duration of the period in which fast dividing neural progenitor cells such as neuroepithelia (NECs) and radial glia cells (RGCs) are active determine both the size and complexity of the neocortex. Neocortical development is characterized by the sequential emergence of distinct neural progenitor subtypes and a concomitant lengthening of the cell cycle. In this study, we demonstrate that transition of the early neural progenitors, specifically RGCs, to intermediate progenitors (IPCs) necessitates loss of expression of the Deltex family member, DTX4, during both human and murine cortical development. We further show that DTX4 maintains RGC identity by shortening cell cycle length. These findings emphasize the importance of cell cycle dynamics for determining progenitor identity and establish DTX4 as a key regulator of these processes and thereby of developmental progression in the neocortex.

### Cell cycle dynamics drive progenitor transitions

Lengthening of the cell cycle length has been proposed as a causal factor in neural progenitor transitions during neocortical development^6,21^. Studies in the murine and ferret neocortex demonstrated that the sequential appearance of different progenitor populations is accompanied by a progressive increase in cell cycle duration^6,22,23^. Furthermore, the manipulation of G1 phase-associated cyclin-dependent kinases in NEC progenitors, altered both cell cycle duration and transition to RGCs ^21,24–27^.

Our findings extend this paradigm to the transition of RGCs to IPCs, underscoring the central role of cell cycle regulation in progenitor fate determination. Specifically, we provide a mechanism for how cell cycle duration is regulated during neural progenitor transition. We show that loss of DTX4 expression is a critical driver of altered cell cycle dynamics and transition to IPCs. Conversely, sustained DTX4 expression shortens the cell cycle and prolongs RGC identity, indicating that DTX4 is both necessary and sufficient to maintain the highly-proliferative RGC state. Elongation of the proliferative period has enabled the evolutionary expansion of the neocortex. It is of note that DTX4 is expressed by highly proliferative progenitor populations, such as oRGCs, that are characteristic of gyrencephalic species^28–31^. An extended period of expression of DTX4 may be involved in enabling cortical expansion and cellular diversification in these species.

## Materials and methods

### Experimental model

#### Mice

All animal experiments were performed at the Charité – Universitätsmedizin Berlin in compliance with the guidelines for the welfare of experimental animals approved by the State Office for Health and Social Affairs, Council in Berlin, Landesamt für Gesundheit und Soziales (LaGeSo), permissions G0055/19, T-CH0033/22, G0184/20, and G0027/24. Wildtype outbred mice (NMRI; Charles River Laboratory) and Satb2-Cre mice^14^ were used in this study. Experimental ages used are indicated in the figure legends and in the main text. Each sample included both sexes within litters without distinctions. Males were single housed from the point of exposure to female animals. Females were housed in groups of 2-4 animals per cage. The date of vaginal plug was counted as E0.5. In adherence with Charité and BIH recommendations, this study used the ARRIVE guidelines 2.0 for reporting animal research. All relevant information can be found in figure legends or in each relevant section. No exclusion criteria were set for this study.

#### Cell lines

One human iPSC line (BIHi250-A, https://hpscreg.eu/cell-line/BIHi250-A) and the murine Neuro-2a neuroblastoma cell line were used in this study.

### Method Details

#### Cerebral organoid generation

Human cerebral organoids were generated from the BIHi250-A line using STEMdiff™ Cerebral Organoid Kit (STEMCELL Technologies #08570 and #08571) according to protocol^32^. Before differentiation BIHi250-A were maintained in E8 media on Geltrex coated plates and differentiated regions removed. hiPSC were incubated in TrypLE Select (Thermo Fisher, 12563029) for 8-10min at 37°C in a 5% CO_2_ incubator. Cells were gently detached, centrifuged and then resuspended in EB Seeding Medium at 90,000 cells/ml and finally plated in 96-well ultralow attachment plates (Costar, 7007) at 9000 cells/well. EB Formation Medium was added on D2 and Induction media on D5. EBs were then transferred at a maximum of four EBs/well to 24-well ultra-low attachment plate and incubated for 48h at 37°C in a 5% CO_2_ incubator. EBs were subsequently embedded in Geltrex and incubated in Expansion Media in an anti-adherent dish at 37°C in a 5% CO_2_ incubator for 3 days to develop an expanded neuroepithelia. The media was replaced with Maturation Media supplemented with 1x Antibiotic-Antimycotic solution (Thermo Fisher, 15240096) on D10 and grown further on an orbital shaker set to 65 rpm at 37°C in a 5% CO_2_ incubator. Maturation media was changed every other day.

### *In utero* electroporation (*IUE*)

*In utero* electroporations were carried out as previously described^15^. Briefly, a DNA mixture (500ng experimental or control plasmid) was prepared in endotoxin-free water with 0.1% Fast Green FCF (Sigma-Aldrich) and injected into the lateral ventricle of E12 embryos using micropipettes prepared from 1.5-1.8 x 100mm borosilicate glass capillaries (Kimble and Chase). DNA uptake by neural progenitors was induced by the application of 6 pulses of 35 across the uterine wall using platinum electrodes.

### *In organoid* electroporation (*IOE*)

*IOE* was performed at D28. The organoids were electroporated one by one. Each organoid was washed once in 1x PBS then moved to a plate with OPTI-MEM (Gibco, 31985-062). The DNA mixture diluted in UltraPure™ DNase/RNase-Free Distilled Water (Invitrogen, 10977049) was injected into three lumens per organoid using a pulled glass capillary (Kimble and Chase) and an Eppendorf CellTram 4r Air Microinjector system. Organoids were electroporated using platinum forceps electrodes (NEPA GENE) and a NEPA GENE Super Electroporator type II using a poring pulse setting of 80 volts (V), 3.5msec pulse length, 50msec pulse interval, 5 pulses, and 10% decay rate and a transfer pulse setting of 20V, 50msec pulse length, 50msec pulse interval, 5 pulses, and 40% decay rate. The organoid was then moved into full COD electroporation media without anti-anti for 24h before moving into full COD media (0.5x DMEM-F12 (Gibco, 31330038), 0.5x Neurobasal (Gibco, 21103-049), 0.5x N2 supplement (Gibco, 21103-049), 1x GlutaMAXTM-I (Gibco, 35050-061), 0.5x MEM Non-Essential Amino Acids (Gibco, 11140035), 2.5 mg/ml Insulin human recombinant (Merck, 91077C-100MG), 45μM 2-Mercaptoethanol (Gibco, 21985023), 0.5x B27 supplement (Gibco, 17504044), 1x Antibiotic-Antimycotic (Anti-anti, Glibco, 15240096)).

### Cell culture and transfection

Neuro-2a mouse neuroblastoma cells were obtained from DSMZ (ACC 148) and grown in Neuro-2a media (50% Dulbecco’s modified Eagle’s medium (DMEM) + GlutaMAX^TM^-1 (Gibco, 31966-021) 50% Neurobasal medium (Gibco, 21103-049) containing 10% fetal calf serum (FCS, Sigma-Aldrich, F9665) and 1% penicillin and streptomycin (Pen Strep) (Gibco, 15140-122)) in a 5% CO_2_ 37°C incubator. Transfection was carried out in OPTI-MEM (Gibco, 31985-062) media using Lipofectamine 2000 (Invitrogen, 11668-019) according to protocol.

### Sample preparation

Embryonic brains were dissected out of the skull and placed in ice cold 4% paraformaldehyde (PFA)/PBS overnight at 4°C. The brains were cryoprotected by incubation through a 15%-30% sucrose/PBS gradient and subsequently frozen in a 2-methylbutane (Roth, 3927.1) dry ice bath. Brains were stored at least overnight at −70°C. Samples were coronally sectioned at 50μm using a RWD Minux FS800 cryostat and placed directly on Superfrost^TM^ Plus glass slides (Epredia, J1800AMNZ). For in situ hybridisation, DEPC was included in all PBS based solutions and samples were sectioned at a thickness of 16μm.

P21 animals were sacrificed by lethal dose of pentobarbital (Narcoren, Boehringer Ingelheim). Mouse unresponsiveness was checked by toe-pinch response. Intracardiac perfusion with 4%PFA/PBS was performed manually. Brains were collected in ice-cold PBS and fixed in 4% PFA/PBS overnight at 4°C. After washing in PBS, brains were embedded on the cutting surface using tissue glue on the base of the brain and 150μm coronal brain sections prepared using a HM650V Microm. Tissue sections were stored in 1x PBS with 0.01% sodium azide at 4°C. After fixation, organoids were cryoprotected by incubation through a 15%-30% sucrose/PBS gradient. Organoids were embedded in Tissue-Tek O.C.T. Compound (Plano, 19255028026) and frozen on dry ice. Samples were stored at −70°C for at least 24h or until sectioning. 16μm sections were prepared using a RWD Minux FS800 cryostat and placed directly on Superfrost^TM^ Plus slides.

### In situ hybridisation

In situ hybridisation was performed on mouse brain cryosections. Sections were post-fixed for 20min with 4% PFA-DEPC (Roth, 1609-47-8) then washed in 1x PBS-DEPC (Roth, 1609-47-8). The tissue was treated with warm (37°C) Proteinase K (20mg/ml, Sigma Aldrich, 1.07393) in Proteinase K buffer (30 mM Tris pH 8.0, 10mM EDTA pH 8, 0.5% SDS) for 2-3min at RT, quickly rinsed in 1x PBS-DEPC and then incubated in 0.2% Glycine in 1x PBS-DEPC for 5min at RT. After washing, slides were incubated in 4% PFA 0.2% Glutaraldehyde (Sigma Aldrich, 340855) in 1x PBS-DEPC for 20min at RT and then re-washed in 1x PBS-DEPC. Samples were then pre-incubated with hybmix (50% Formamide (Sigma Aldrich, F9037), 5x SSC (Sigma Aldrich, S6639), 1% Boehringer block (Sigma Aldrich, C7897), 5mM EDTA, 0.1% TWEEN® 20 (Sigma Aldrich, P1379), 0.1% CHAPS (Roche, CHAPS-RO), 0.1mg/ml Heparin (Sigma Aldrich, H0878), 100 μg/ml Yeast RNA (Roche, 10109223001) in DEPC-Water) at 68°C for 2h before incubation with the denatured probe diluted in hybmix at 68°C overnight. Slides were washed once in 2XSSC pH4.5 and then incubated three times for 30min at 65°C in wash solution (50% Formamide/2X SSC pH 4.5) and once with KTBT (50mM Tris pH 7.5, 150mM NaCl, 10mM KCl, 1% Trition-X100) for 10min at RT. The slides were then incubated for 2h at RT in blocking solution (20% sheep serum (Sigma Aldrich, S2263) in KTBT), followed by overnight incubation at 4°C with anti-DIG alkaline phosphatase antibody (Sigma Aldrich, A3076) diluted 1:1000 in blocking solution. The slides were thoroughly washed in KTBT 5min followed by one wash in 2x in NT(M)T (100mM Tris pH 9.5, 100mM NaCl, 50mM MgCl2, 0.05% TWEEN® 20) all at RT. Staining was carried out by incubation in staining solution (1:50 NBT/BCIP (Roche, 11681451001) in NT(M)T) until colour developed. After washing in 1x PBST (PBS + 0.1% Triton X100), slides were then incubated in 4% PFA for 10min at RT and mounted in 40°C Organic rapid mounting medium (Sigma Aldrich, 1.07961). Samples were imaged on an AxioVision BX60 microscope.

### Immunofluorescence

All samples of an experiment were stained in parallel on samples derived from littermate animals. Sections were first checked for GFP signal using Axiovert 40 CFL microscope. Only GFP^+^ sections were stained. Brain sections of P21 animals were first incubated for 5 minutes with 1mg/ml sodium borohydride (Sigma-Aldrich, 213462). Embryonic, postnatal and organoid samples were then washed in 1x PBS before incubation in blocking solution (10% horse serum, 0.5% Triton X-100 in PBS) for 1h at RT. Incubation with primary antibodies and nuclear stain (DRAQ5, Invitrogen or DAPI, Sigma-Aldrich) diluted in blocking solution was carried out overnight at 4°C. After three washes in 1x PBS, samples were incubated in a 1:300 dilution of secondary antibodies in blocking solution for 4-6h at RT, washed three times in 1x PBS and mounted on Superfrost^TM^ Plus glass slides using a drop of Immu-Mount (Epredia, 9990402) and a coverslip (Roth, 24’50/2424’60, 1871). Zeiss Yokogawa Spinning Disc CSU-X1, Nikon CSU-W1 SoRa and Nikon Confocal AX-NSPARC microscopes were used to image the samples. All samples of an experiment were imaged using the same laser intensity and settings.

### EdU-BrdU double labelling

One day after transfection, cells were labelled by addition of 10mM EdU for 1.5h followed by addition of 10mM BrdU for an additional 30min incubation at 37°C 5% CO_2_. The cells were fixed in 4% PFA 1x PBS at RT before staining.

### Hoechst staining and Flow cytometry

24h after transfection, cells were washed in 37°C PBS. The washing solution was removed, and the cells were incubated with 5μg/ml Hoechst (Invitrogen, H1399) in Neuro-2a media without anitbiotics for 30min at 37°C 5% CO_2_ incubator. The staining solution was removed, and the cells were incubated with 0.25% trypsin without EDTA (Gibco, 15090-046) with 5μg/ml Hoechst (Invitrogen, H1399) for 2min at 37°C 5% CO_2_ incubator. Cells were then dissociated in PBS with 1% FCS (Sigma-Aldrich, F9665). The cell suspensions were then sorted using BD FACSCanto II according to their GFP and Hoechst profiles to determine the cell cycle profiles of each condition.

### Quantification and Statistical analysis

#### Quantification of distribution of cortical neurons

The laminar distribution of modified cells and cell fate analysis were performed as described previously^15^. Briefly, Z stack of 50μm thick *in utero* electroporated murine cortices or 16μm sectioned human cerebral organoids were acquired with Zeiss -spinning disc CSU-X1,Yokogava, Axio Observer Z1 (murine brain cortices) or Nikon CSU-W1 SoRa (organoids). The position of each modified neuron was marked using the Cell Counter plug-in of ImageJ software and plotted with respect to the position of the ventricular surface or organoid lumen surface. For analysis of the distribution of different cell fate or cell cycle markers, positive cells were identified by eye then quantified using the Cell Counter plugin of Image J and represented as percentage of all GFP^+^ cells in the section.

#### Quantification of crossing callosal axons

Mean GFP intensity of callosal axons was measured at the midline and normalized to the GFP intensity at the site of *IUE* using the Multi Measure plugin in ImageJ. The same ROI was used for all brains.

#### Determination of cell cycle length

Cell cycle length was calculated as previously described^20^, using the following equations Ts = Ti / (L cells / S cells), L cells = EdU^+^BrdU^-^, S cells = EdU^+^BrdU^+^. Ts was then used to calculate cell cycle length using Tc = Ts / (S cells / P cells) where S cells = EdU^+^BrdU^+^ and P cells = number of proliferative cells determined by counterstaining with Ki67 and GFP.

#### Cell cycle profiles

Flow cytometry was carried out using the BD FACSCanto II, Gating parameters for GFP and Hoechst signal, were set according to published protocols^33^. The online software Floreada.io (https://floreada.io/analysis) was used to analyse the flow cytometry data. The signal following the gating was then used for cell cycle analysis using the built-in Floreada.io Cell Cycle function.

#### Invagination index

The invagination index was calculated as previously described^34^. In brief, the contour of the cortical surface of layer 1 at the site of invagination and the contour of the overlying pia was outlined using the same starting and ending positions using the outline tool in ImageJ. The length of both lines was measured using the Multi Measure plugin in ImageJ. The Invagination Index was calculated by dividing the length of the layer 1 contour by the length of the pia contour.

#### Data analysis

Analysis of immunofluorescent imaging was done using Fiji Image J. Graphs and statistics were prepared in Prism GraphPad 10. The assumption of normality of parametric analysis were tested using Kolmogorov-Smirnov and Shapiro-Wilk. Homoscedasticity was assessed using plots generated in GraphPad Prism 10. Non-parametric testing was done where appropriate.

Type 1 (false positives) and 2 errors cannot be completely avoided. However, they were minimized by using an appropriately large sample size and statistical power which was calculate a-priori using the statistical software G*Power (version 3.1.9.7)^35^.

## Acknowledgements

This work was supported by the German Research Foundation (DFG) grants RO 3497/3 and RO 3497/4 (M.R.) and the Einstein Center for Neurosciences Berlin (E.P.). We would like to acknowledge the technical support of the BIH Cytometry Core Facility. In addition, we thank Jutta Schueler for assistance with microscopy.

## Author contributions

E.P. conducted and quantified the majority of the experiments. S.Z. aided in the in vivo analysis.

X.D. carried out the *in situ* hybridizations. E.E. carried out preliminary *IUE*s to P21. P.M.W. processed preliminary *IUEs* to P21. D.L. provided technical assistance throughout the project.

T.S. cloned DTX4. M.C.A. contributed feedback. Human cerebral organoids were generated by

V.F.V. and maintained by E.P. under the administration of H.S.. Samples for LC-MS were prepared by E.P., M.R. and D. L.. LC-MS was carried out by K.T.T.. Experiments were analysed by E.P. and M.R.. M.R. conceptualized, conceived and supervised the study. The manuscript was written and figures were prepared by E.P. and M.R.. The manuscript was reviewed by all authors.

