## Supplementary Information for "DTX4 regulates neural progenitor transitions during cortical development"

### Supplementary Methods

#### Preparation of cell lysates

Cells were harvest 24h after plating or 24h after transfection in RIPA buffer (50mM Tris pH 8.0, 150mM NaCl, 0.1% SDS, 1% TERGITOL, 0.5% Sodium Deoxycholate) with inhibitors (1x PhosStop (Roche, 04906837001), 1x Protease inhibitor cocktail (PIC) (Roche, 11836170001), 5µg/ml Pepstatin (Sigma Aldrich, P5318), 2.5mM Na vanadate (Sigma Aldrich, S6508), 5mM NaF (Santa Cruz, sc-24988A), 10mM Benzamidine (SAFC, 06837), 10 µg/ml Leupeptin (Sigma Aldrich, L2884), 5µg/ml Aprotinin (Thermo Scientific™, 78432), 1x MG132 (Sigma Aldrich, 474790), 10mM NEM (Sigma Aldrich, 04260), 0.2mM PMSF (Sigma Aldrich, P7626), 1:1000 Benzonase (SAFC, 1.01654), 2mM $MgCl_2$). The lysates were cleared by centrifuged at 13 500 rpm for 10 min at 4°C (Eppendorf centrifuge 5427-R). Protein concentrations were measured by Bradford assay using Bio-Rad Protein Assay reagents. Samples prepared in 1x Laemmli buffer (25mM Tris, 192mM glycine and 0.1% SDS) for SDS-PAGE.

#### SDS-PAGE, Western blotting

Samples were subjected to SDS-PAGE and resolved proteins were then transferred to PVDF membranes for Western blotting. Briefly, transfer to PVDF membranes was carried out in transfer buffer (25 mM Tris pH 7.4, 192 mM glycine, 10% methanol) at 100V for 90min at 4C using a wet transfer setup (Bio-Rad Western Blotting Equipment). Membranes were washed in TBST (10mM Tris pH 8.0, 150mM NaCl, 0.05% TWEEN® 20), blocked in 3% BSA/TBST and then incubated overnight in primary antibody diluted in 1%BSA/TBST. Following washes, membranes were incubated in secondary antibodies in 1%BSA/TBST for 2h at RT. Signal was developed using 500ml SuperSignal$^{TM}$ West Pico PLUS Chemilumiescent Substrates (Thermo Scientific, 34580) and detected on an Azure Biosystems 600 Imager. Western blot images were quantified using Azure Biosystems Western Blot Analysis Software and Prism GraphPad 10 was used for data presentation and analysis.

#### Preparation for primary cortical neurons

Satb2$^{Cre/+}$ E12 embryos cortices were prepared in ice cold 1x Hanks' Balanced Salt Solution (HBSS, +$CaCl_2$, +$MgCl_2$) (Gibco, 14025-050) without phenol red. The cortices were washed in ice cold 5ml HBSS (-$CaCl_2$, -$MgCl_2$, Gibco, 14170-088) with phenol red and cells were then dissociated by incubation in 0.3125% trypsin solution at 37°C for 15min with gentle mixing after

5-7min. After trypsin incubation, the cortices were centrifuged at 900rpm for 5min at 20°C (Eppendorf centrifuge 5427-R). The supernatant was carefully removed, and the cortices were washed in plating media (DMEM + 10% FSC). DNase (Roche, 04716728001) was added for 2min at 37°C to remove free DNA. Cells were collected by brief centrifugation, washed once in plating media and resuspended in plating media.

#### Nucleofection of $Satb2^{Cre/+}$ NMRI E12 primary neurons

Primary cortical neuron suspensions were nucleofected using the Mouse Neuron Nucleofector kit (VPG-1001, Amaxa 2b Nucleofection system, Lonza) and program O-005 on nucleofector (Lonza) following manufactures instructions. Specifically, $5\times10^6$-$2\times10^7$ cells mixed with DNA at a concentration of 1$\mu$g DNA per $1x10^6$ cells. After nucleofection, cells were plated and incubated for 24h at 37°C 5% $CO_2$.

#### Flow cytometry of primary cortical neurons

24h after nucleofections, cells were washed with 37°C HBSS +$CaCl_2$, +$MgCl_2$ (Gibco, 14025-050) and then collected by trypsinization in 0.25% trypsin without EDTA (Gibco, 15090-046) for 3min in a 37°C 5% $CO_2$ incubator. Trypsinization was ended by resuspension in 1% FCS/PBS. The cells were sorted according to their Red/Green fluorescent profiles with the assistance of the BIH Cytometry Core Facility using a BD FACSSymphony A3 (BD Biosciences, San Jose, CA, USA), configured with 5 lasers (UV, violet, blue, yellow-green, red) at the BIH Cytometry Core Facility, with BD FACSDiva Software Version 9.3.1.

#### scRNA data

Murine scRNA data was from Di Bella et al (2021) and cRNA data from human organoids was from Uzquiano et al (2022) and were accessed through https://singlecell.broadinstitute.org/single_cell/study/SCP1290/molecular-logic-of-cellular-diversification-in-the-mammalian-cerebral-cortex#study-summary and https://singlecell.broadinstitute.org/single_cell/study/SCP1756/10.1016/#study-summary respectively.

### Supplementary Figures

Pedersen et al.

Supplementary Fig. S1

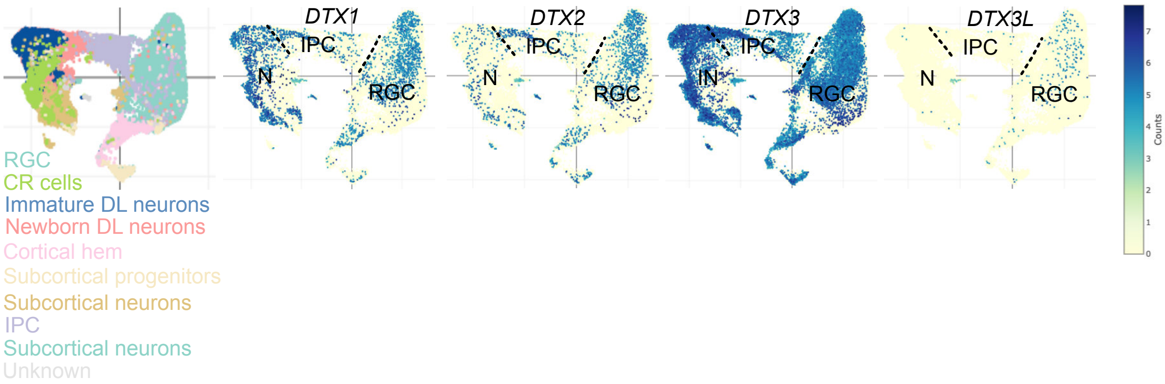


**Fig. S1. DTX4 is expressed in human neocortical progenitor cells.** scRNAseq data from D30 human cerebral organoids from Uzquiano et al (2022). UMAP representation of gene expression profiles for DTX family members: *DTX1, DTX2, DTX3, DTX3L*. RGC, radial glial cell; IPC, intermediate progenitors; N, immature and mature neurons.

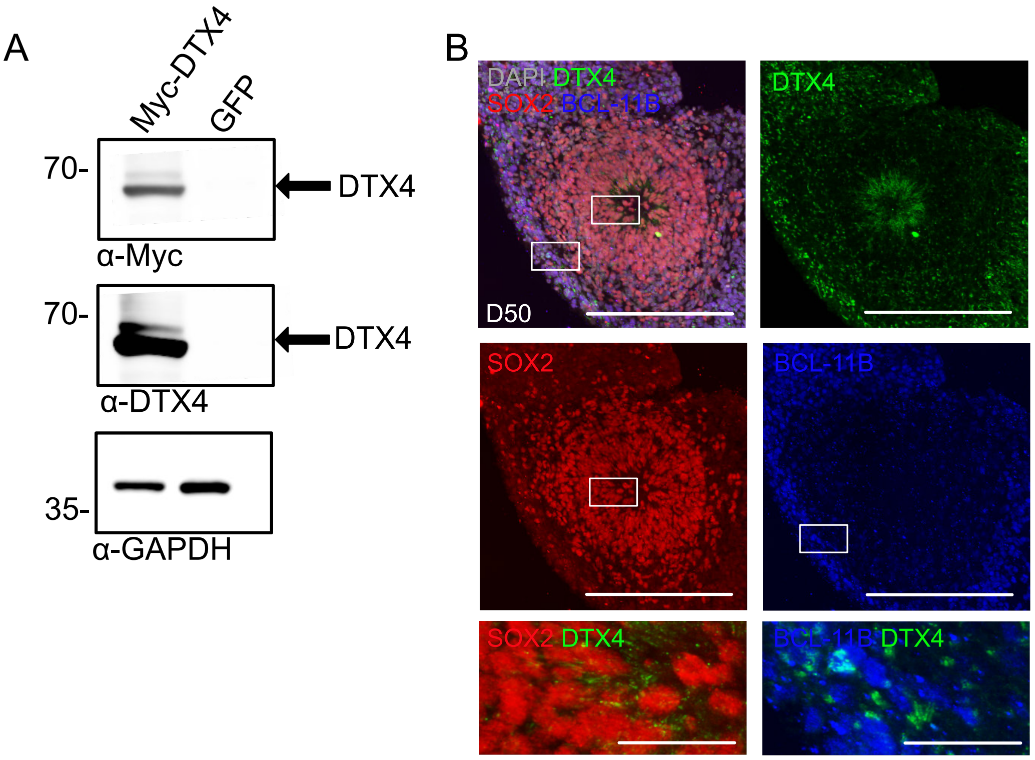


**Fig. S2. DTX4 is expressed in human cerebral organoids.** (A) DTX4 antibody is specific. Western blotting for Myc, DTX4 and GAPDH of Neuro-2a lysates expressing Myc-tagged DTX4 or GFP. (B) DTX4 is expressed in DL neurons in human cerebral organoids at D50. Immunofluorescent staining of D50 WT BIH-250-human cell line cerebral organoid for DTX4 (green), SOX2 (red), BCL-11B (blue), and DAPI (gray) is shown Scale bar = 200μm, magnification 50μm.

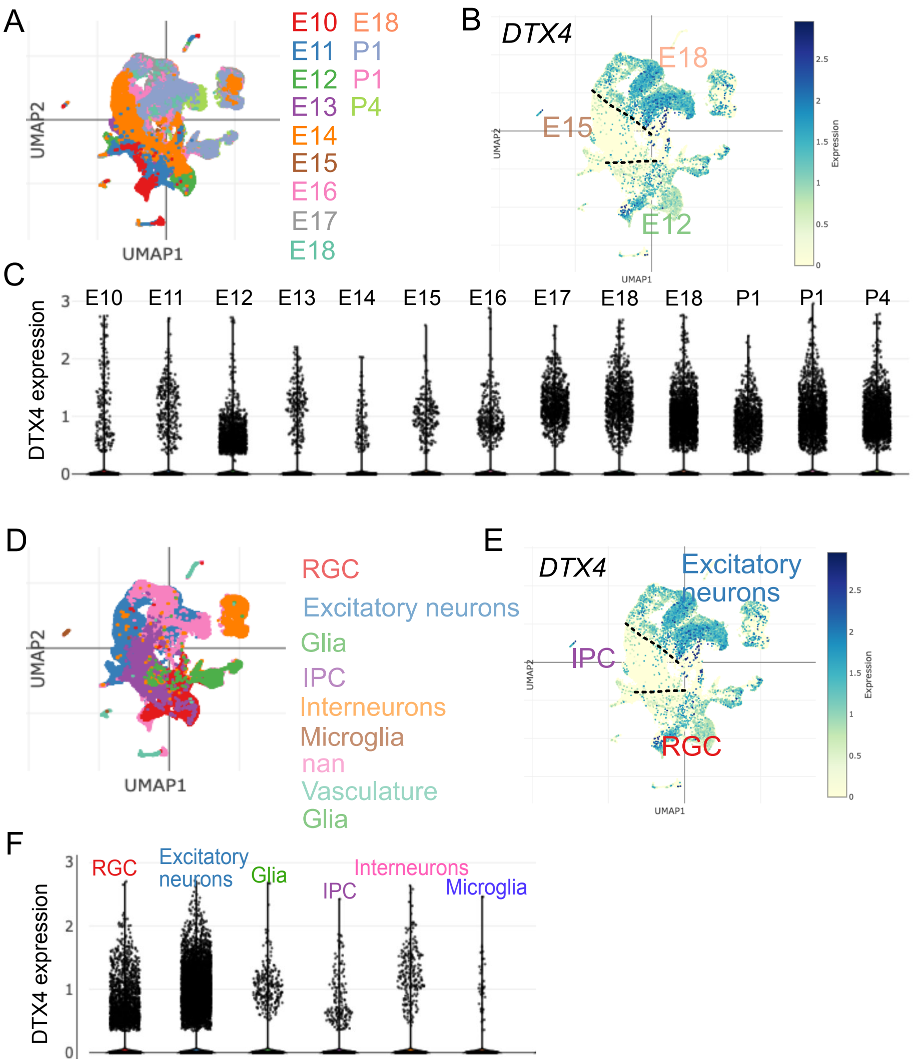


**Fig. S3. DTX4 is expressed at different stages of murine neocortical development**. (A-F) DTX4 is highly expressed at E12 and E18 but not at E15 during murine corticogenesis. (A) UMAP of age clustering of murine neocortex scRNAseq results from Di Bella et al (2021). (B) UMAP of DTX4 expression (C) Graph showing DTX4 expression across developmental age during murine neocortical development. (D) UMAP of cell type clustering from Di Bella et al (2021). (E) UMAP of DTX4 expression showing major cell type clusters (F) DTX4 expression across the different cell type clusters.

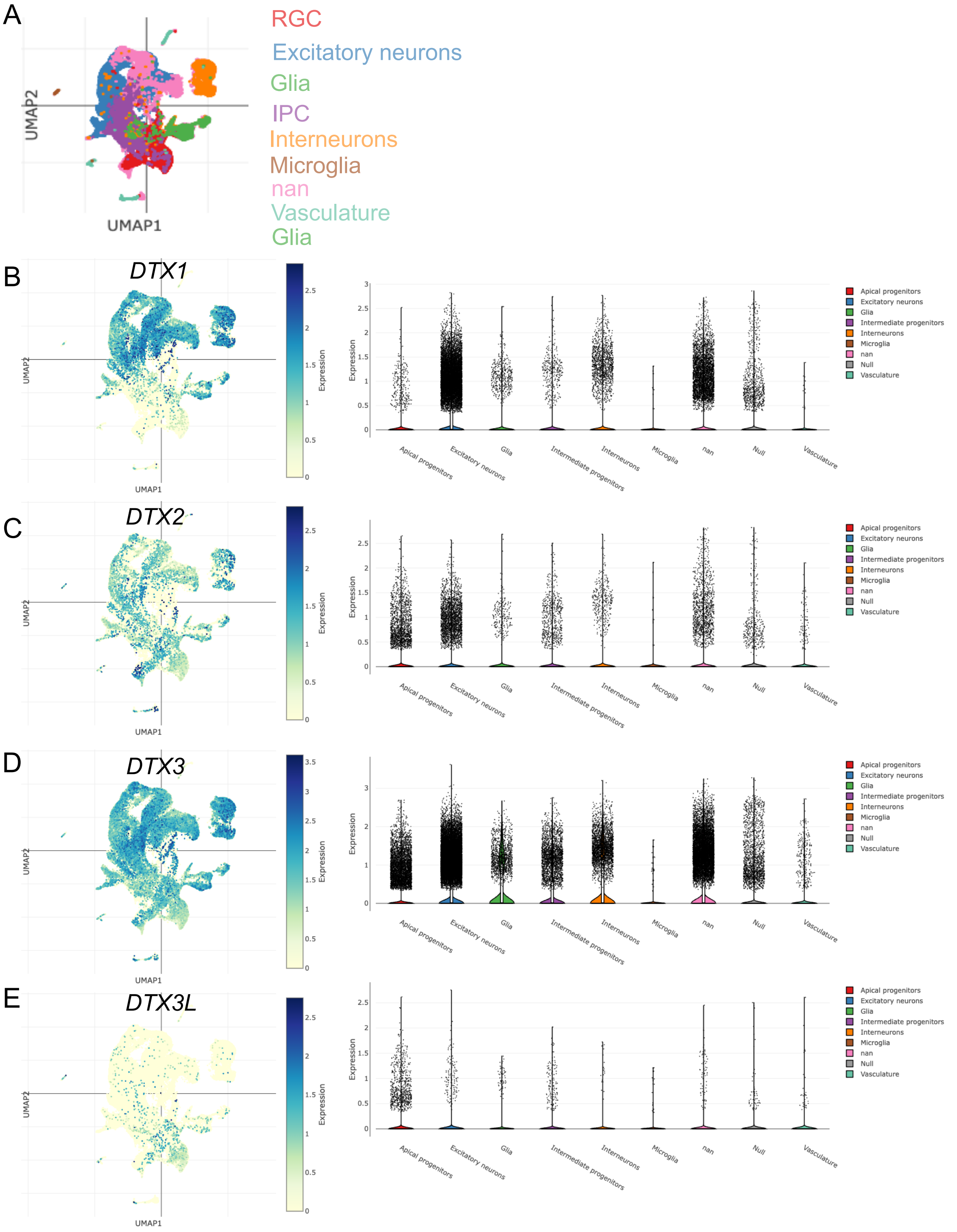


**Fig. S4. Other Deltex family members do not show a biphasic expression pattern during murine corticogenesis.** (A) UMAP of cell type clustering of murine neocortex scRNAseq results from Di Bella et al (2021). UMAP for (B) DTX1, (C) DTX2, (D) DTX3 and (E) DTX3L expression including distribution among cell type clusters.

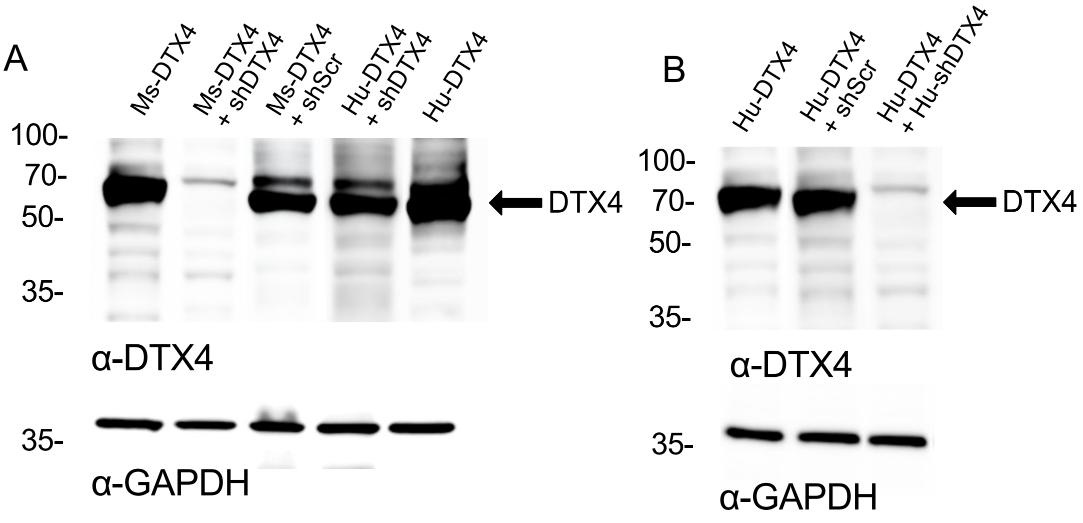


**Fig. S5. Validation of specific shRNAs against (A) murine and (B) human DTX4.** Neuro-2a cells were transfected with either murine DTX4 (Ms-DTX4), human DTX4 (Hu-DTX4), control shRNA (shScr) or an shRNA specific against either murine DTX4 (A) or human DTX4 (B; Hu-shDTX4). Western blotting for DTX4 and for GAPDH are shown.

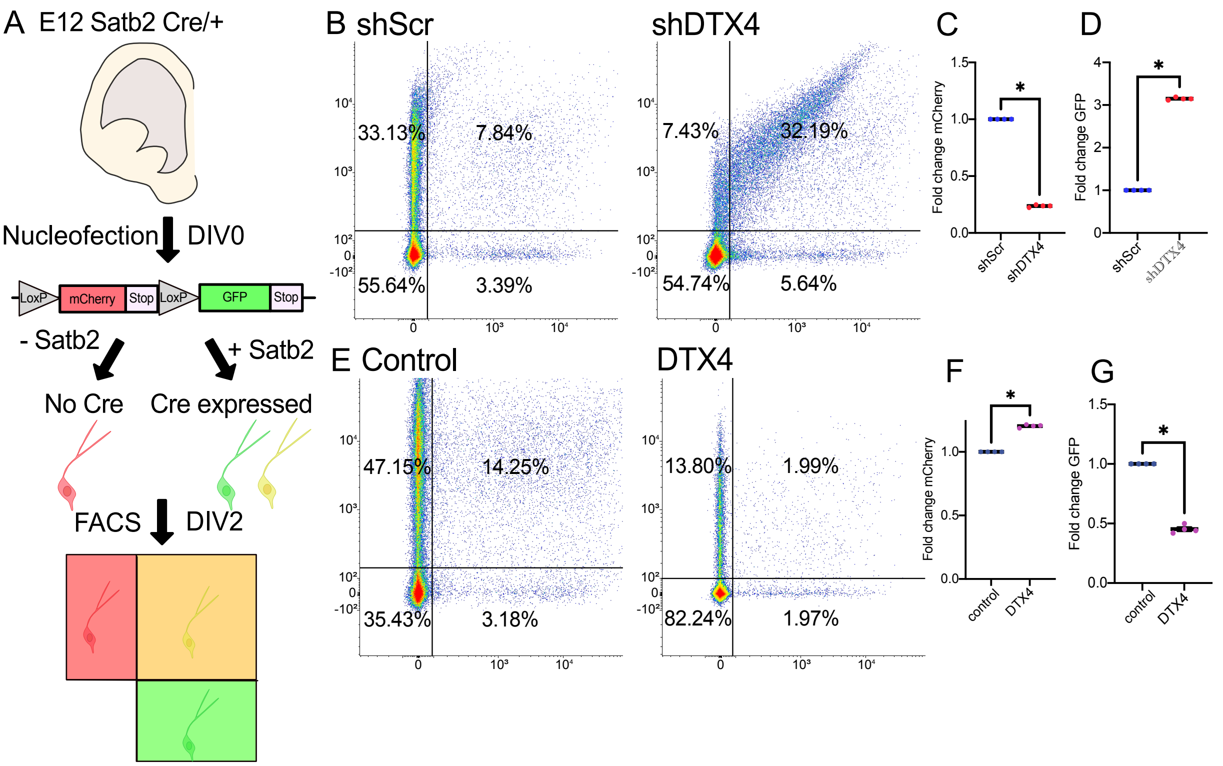


**Fig. S6. DTX4 regulates UL neurogenesis.** (A) Schematic of experimental setup. E12 Satb2Cre+ mouse cortices was dissociated, and the cells were nucleofected with a floxed mCherry-GFP reporter construct, followed by identification of GFP (510-530nm) and mCherry (600-650nm) fluorescence by flow cytometry (B) Representative examples of mCherry/GFP profiles for shScr and shDTX4 nucleofected cells. (C) Fold change in the proportion of mCherry$^{+}$ cells between shScr and shDTX4 nucleofected cells. Non-parametric analysis using Kolmogorov-Smirnov test, p= 0.0286. (D) Fold change in the proportion of GFP+ cells between shScr and shDTX4 conditions. Non-parametric analysis using Kolmogorov-Smirnov test, p= 0.0286. (E) Representative examples of mCherry/GFP profiles for control (empty vector) and DTX4 overexpressing cells. (F) Fold change in the proportion of mCherry+ cells between control and DTX4 conditions. Non-parametric analysis using Kolmogorov-Smirnov test, p= 0.0286. (G) Fold change in the proportion of GFP+ cells between control and DTX4 conditions. Non-parametric analysis using Kolmogorov-Smirnov test, p= 0.0286. All data presented as individual points, mean and error bars with SEM. N=4 (500 000-1 000 000 cells per N). Normality of the data was evaluated using Shapiro-Wilk or Kolmogorov-Smirnov test. BD FACSDiva Software Version 9.3.1 and Floreada.io was used in the processing of the FACS data.

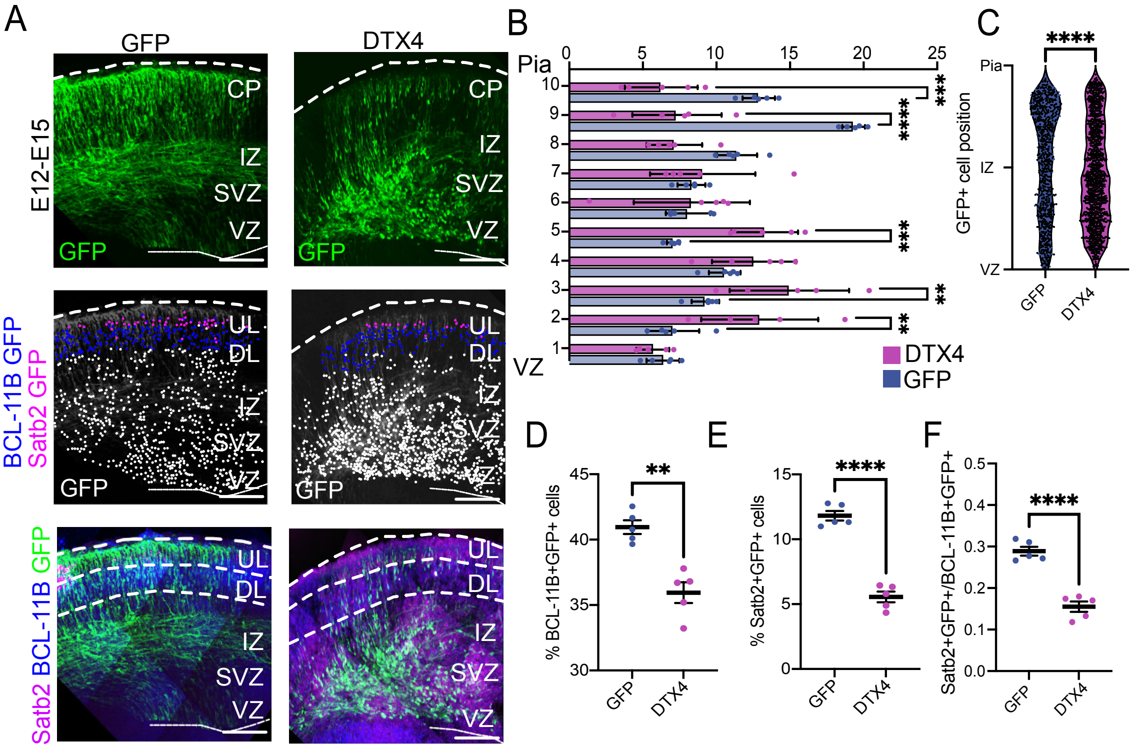


**Fig. S7. DTX4 OE delays neurogenesis in the murine cortex**. (A) Representative examples of brains electroporated at E12 and analyzed at E15 for GFP control and DTX4 OE. The sections were stained for GFP (green, electroporated cells), BCL-11B (blue) and Satb2 (magenta) as indicated. The position of single $GFP^+$ (white), $BCL\text{-}11B^+GFP^+$ (blue) and $Satb2^+GFP^+$ (magenta) cells is marked by colored dots in the middle panels. Scale bar = 100μm (B) Bin graph showing normalized distribution of $GFP^+$ cells across the cortex in 10 bins. 2-way ANOVA analysis. N = 5. (C) Laminar distribution of $GFP^+$ cells in the cortex. Data was analyzed using the non-parametric Kolmogorov-Smirnov test, N=1378 (GFP) and 1167 (DTX4) from 5 embryos each. (D) Percentage of $BCL\text{-}11B^+$ $GFP^+$ cells. Data was analyzed using a two-tailed Welch's T-test. N = 5. (E) Percentage of $Satb2^+GFP^+$ cells. Data was analyzed using a two-tailed Welch's T-test. (F) Ratio of $Satb2^+GFP^+$ / $BCL\text{-}11B^+$ $GFP^+$ cells. Data was analyzed using a two-tailed Welch's T-test. N = 5. All data presented as individual points, mean and error bars with SEM.

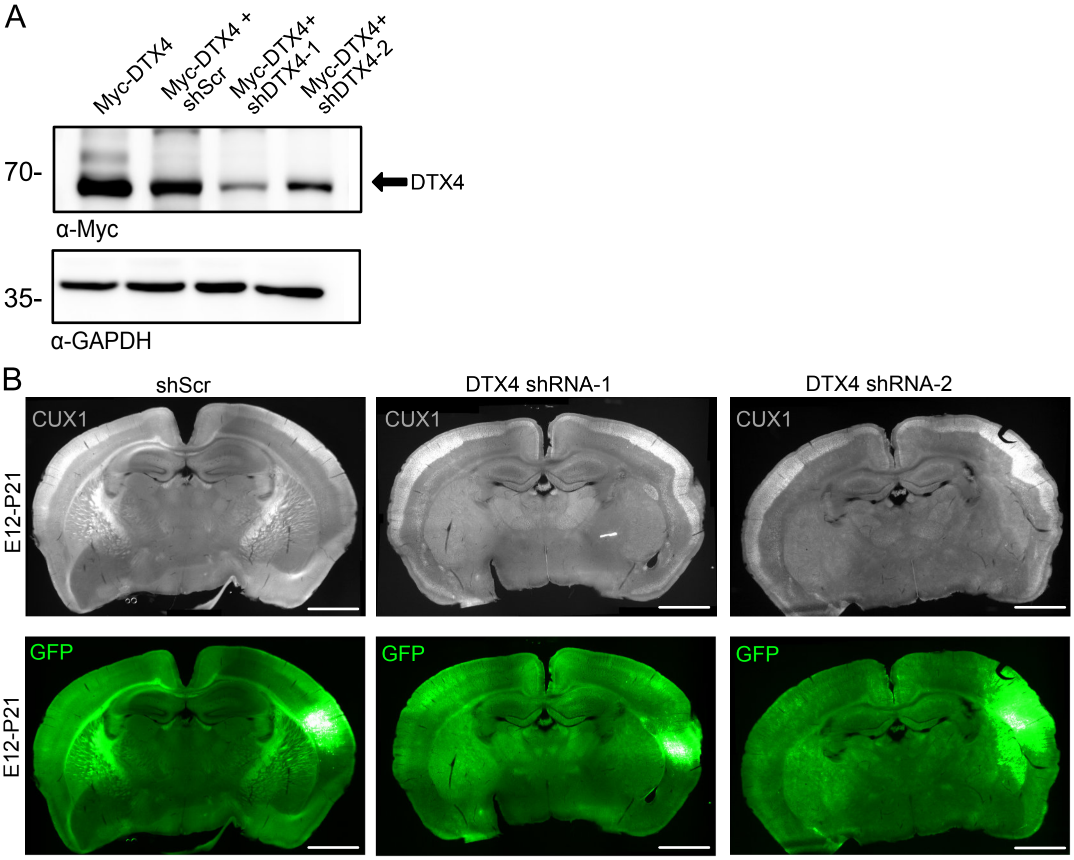


**Fig. S8. DTX4 KD P21 phenotype can be verified using a second specific shRNA for DTX4**
(A) shRNA validation in Neuro-2a cells transfected with Myc-tagged DTX4 and either shScr, shDTX4-1, or a second shRNA for DTX4, shDTX4-2. Western blotting for Myc and GAPDH are shown. (B) Representative images of P21 mouse brains *IUE* at E12 with shScr, shDTX4-1 or shDTX4-2 constructs and GFP then stained with GFP (green) and the UL neuronal marker CUX1 (grayscale). N= 8 shScr, 10 shDTX4-1, and 5 shDTX4-2. Scale bar = 500μm

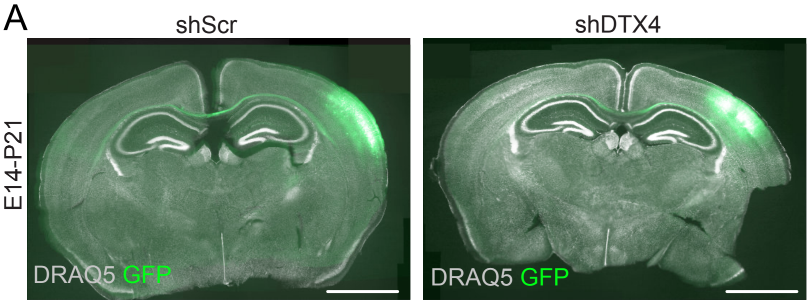


B

| Conditions | Number of brains with cortical malformation | Number of brains without cortical malformation | Percentage of brains with cortical malformation |
|---|---|---|---|
| shDTX4 | 0 | 4 | 0% |
| shScr | 0 | 4 | 0% |

**Fig. S9. DTX4 does not alter cortical morphology during late development**.

(A) Representative images of P21 mouse brains electroporated at E14 with GFP and either shScr or shDTX4. Staining for GFP (green) and the nuclear marker DRAQ5 (greyscale) are shown. Scale bar = 500μm. (B) Overview of the number of brains analyzed per condition and their morphology.

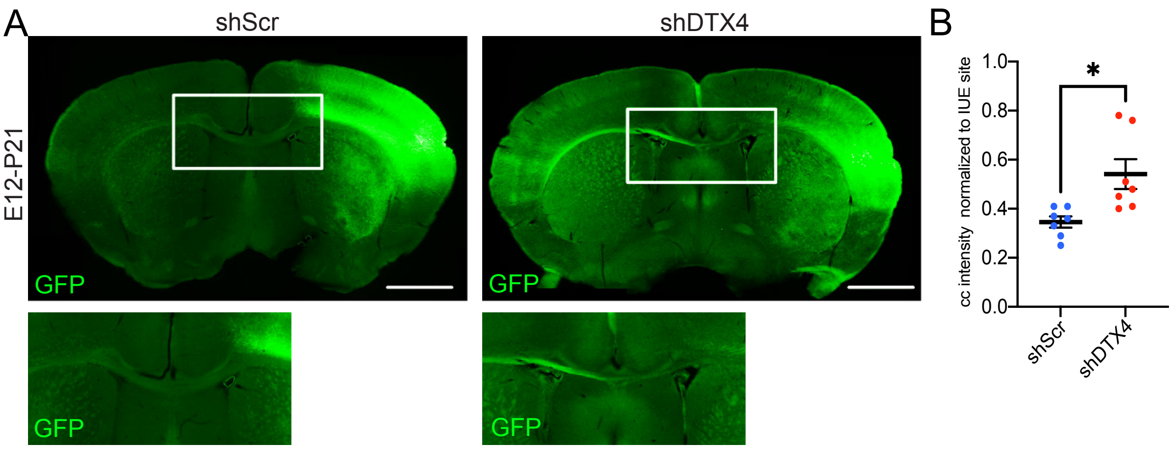


**Fig S10. DTX4 KD increases corpus callosum projecting neurons at P21.** (A) Representative examples of P21 mice *IUE* at E12 with GFP and either shScr or shDTX4 showing magnifications of the corpus callosum. Staining for GFP (green) is shown. Scale bar = 200µm. (B) Normalized corpus callosum (cc) intensity at the midline. Normality of data was evaluated using a Kolmogorov-Smirnov and Shapiro-Wilk tests then analyzed using a two tailed Welch's t-test, N = 7. All data presented as individual points, mean and error bars with SEM.

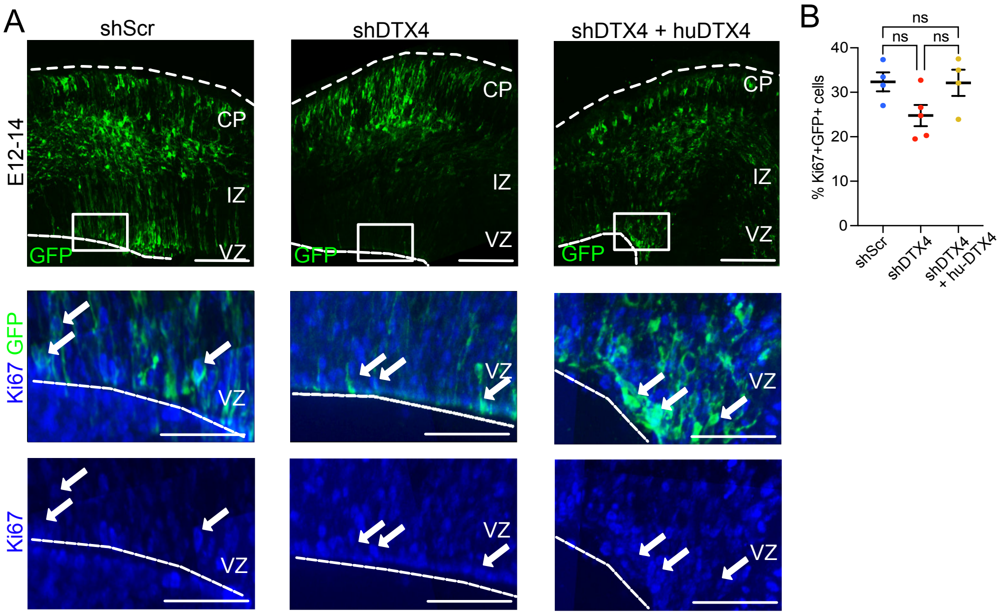


**Fig. S11. DTX4 KD does not alter the number of proliferating cells in the mouse neocortex.** (A) Representative images of shScr, shDTX4, mu-shDTX4+hu-DTX4 (rescue) E12 electroporated brains at E14. Staining for GFP (green) and Ki67 (blue; proliferating cells) is shown. Scale bar = 100μm. Double positive cells are indicated with filled white arrows in the magnified images, scalebar = 50μm. (B) Percentage of $Ki67^{+}$ $GFP^{+}$ cells analyzed by one-way ANOVA with Dunn's correction, N = 5 and 4 for shScr and shDTX4+hu-DTX4 respectively.

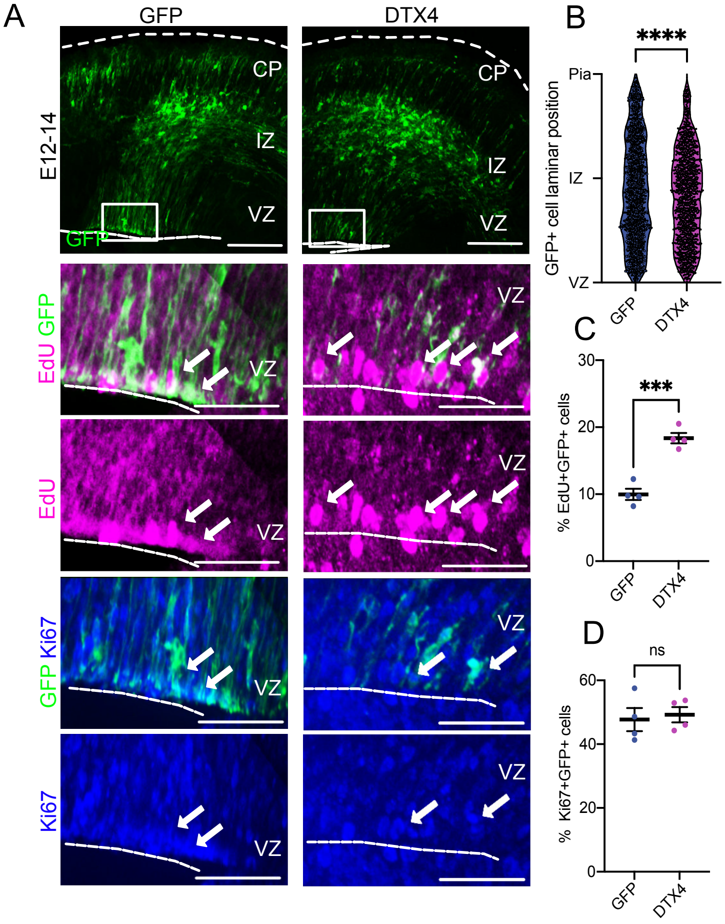


**Fig. S12. DTX4 OE increases the number of S-phase cells but not the number of proliferating cells**. (A) Representative images of mouse brains *IUE* at E12 with either GFP alone or with an DTX4 OE construct. Embryos were exposed to a 2h EdU pulse 36h after *IUE* and analyzed at E14 for expression of GFP (green), incorporation of EdU (magenta) and for expression of Ki67 (blue; proliferating cells). Scale bar = 100μm. Magnified regions are shown by boxes. Double positive cells are indicated with filled white arrows in the magnified images, magnification scalebar = 50μm. (B) Laminar distribution of $GFP^+$ cells in the cortex. Data was analyzed using the non-parametric Kolmogorov-Smirnov test, N= 1416 (GFP) and 1335 (DTX4) from 4 embryos per condition. (C) Proportion of $EdU^+GFP^+$ cells analyzed using an unpaired two-tailed Welch's T-Test, N=4. (D) Proportion of $Ki67^+GFP^+$ cells analyzed using an unpaired two-tailed Welch's T-Test, N=4. Normality of the data was evaluated using Shapiro-Wilk or Kolmogorov-Smirnov test. All data presented as individual points, mean and error bars with SEM except for the box plot which shows individual data points, median, and quartiles.

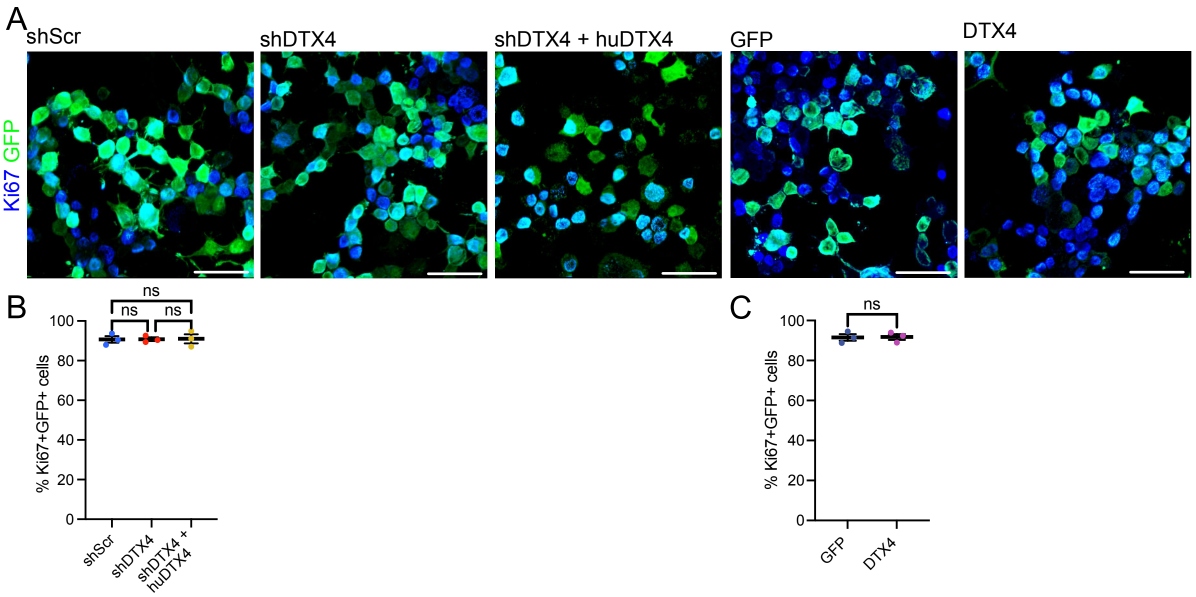


**Fig. S13. DTX4 does not alter overall number of proliferating cells.** (A) Representative images of Neuro2a cells transfected with shScr, shDTX4, shDTX4+huDTX4, GFP, or DTX4 as indicated then stained for GFP (green) and Ki67 (blue). Scale bar = 50μm. (B) Proportion of $Ki67^{+}GFP^{+}$ cells for all conditions. One-way ANOVA with Dunn's correction. N=3 independent transfections per condition (shScr N1= 456, N2= 415, N3= 304, shDTX4 N1= 406, N2= 544, N3= 491, shDTX4+hu-DTX4 N1= 405, N2= 534, N3= 293). (C) Proportion of Ki67+GFP+ cells after transfection with GFP or DTX4. Unpaired two-tailed Welch's T-Test, N=3 independent transfections per condition (GFP N1= 401, N2= 475, N3= 402, DTX4 N1= 511, N2= 299, N3= 408). Normality of the data was evaluated using Shapiro-Wilk or Kolmogorov-Smirnov test. All data presented as individual points, mean and error bars with SEM.

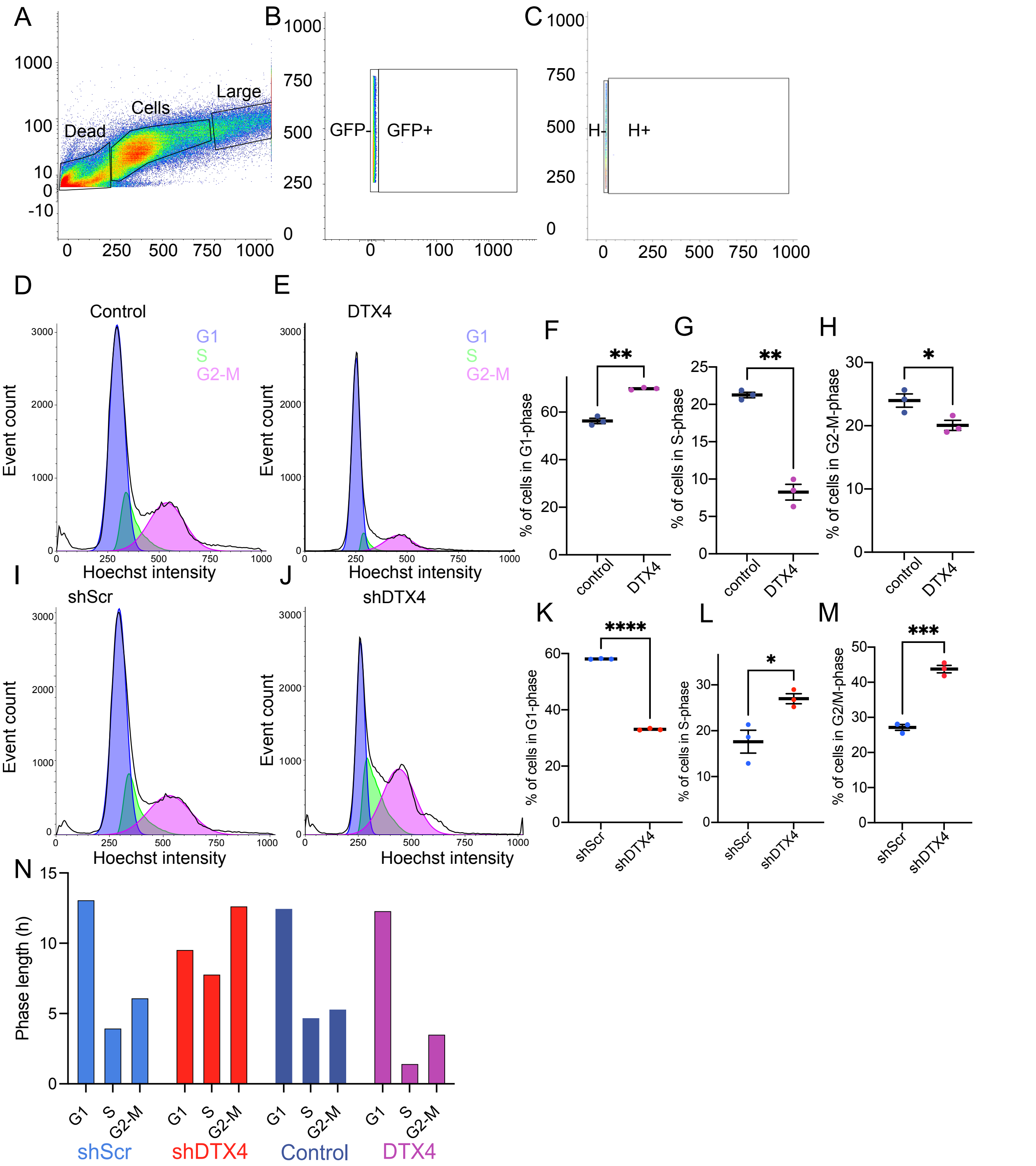


**Fig. S14. DTX4 regulates cell cycle dynamics.** (A) Flow cytometry gating parameters. SSC (y-axis), FSC (x-axis). (B) GFP gating of non-transfected, non-treated control group using FSC (y-axis) and GFP signal (x-axis). (C) Hoechst gating of non-transfected, non-treated control. group showing using FSC (y-axis) and Hoechst signal (x-axis). (D-H) DTX4 OE increases the proportion of cells in G1. Representative cell cycle profiles for cells transfected with either (D) an empty vector (control) or (E) an DTX4 overexpression construct. Proportion of cells in (F) G1, (G) S, and (H) G2/M phase for the different conditions. The data was analyzed using a two-tailed Welch's T-Test. (I-M) DTX4 downregulation results increased the proportion of cells in S and G2-M phase of the cell cycle. Representative cell cycle profile for shScr control (I) and shDTX4 (J) transfected cells are shown. The proportion of cells in (K) G1, (L) S and (M) G2/M-phase are shown for each condition. The data was analyzed using a two-tailed Welch's T-Test. (N) Predicted cell cycle phase length for shScr, shDTX4, EV and DTX4. Calculations are based on percentage of cells in each phase and the overall cell cycle lengths established in Fig. 5 and Supplementary Fig. S13. N=3 (500 000-1 000 000 cells per N). Normality of the data was evaluated using Shapiro-Wilk or Kolmogorov-Smirnov test. All FACS data was analyzed using the online software Floreada.io (https://floreada.io/analysis). All data are presented as individual points, mean and error bars with SEM
